# Quantitative imaging of chromatin decompaction in living cells

**DOI:** 10.1101/219253

**Authors:** Elisa Dultz, Roberta Mancini, Guido Polles, Pascal Vallotton, Frank Alber, Karsten Weis

## Abstract

Chromatin organization is highly dynamic and regulates transcription. Upon transcriptional activation, chromatin is remodeled and referred to as “open”, but quantitative and dynamic data of this decompaction process are lacking. Here, we have developed a quantitative high-resolution microscopy assay in living yeast cells to visualize and quantify chromatin dynamics using the *GAL7-10-1* locus as a model system. Upon transcriptional activation of these three clustered genes, we detect an increase of the mean distance across this locus by >100 nm. This decompaction is linked to active transcription but is not sensitive to the histone deacetylase inhibitor trichostatin A or to deletion of the histone acetyl transferase Gcn5. By contrast, the deletion of *SNF2* (encoding the ATPase of the SWI/SNF chromatin remodeling complex) or the deactivation of the histone chaperone complex FACT lead to a strongly reduced decompaction without significant effects on transcriptional induction. Our findings are consistent with nucleosome remodeling and eviction activities being major contributors to chromatin reorganization during transcription but also suggest that transcription can occur in the absence of detectable decompaction.

## Introduction

DNA in the cell nucleus is present as chromatin in a tight complex with histones and other proteins. This complex is central to spatially organize the DNA strand by balancing the negative charges of the phosphate backbone and is also crucial for gene regulation. The three-dimensional chromatin conformation is highly dynamic and remodeled continuously as cells change their physiological state or their transcriptional programs. This remodeling is orchestrated by histone modifiers and chromatin remodelers, which change the interaction between nucleosomes as well as the interaction between histones, DNA and the protein complement present at a chromatin site, thereby affecting the spatial packing of nucleosomes, their location, mobility or density.

Activation of transcription typically leads to a change in chromatin conformation manifested in higher accessibility of the DNA to digestion or transposon integration (Tsompana and Buck, 2014). Although the changes in histone occupancy and accessibility have been studied extensively, the quantitative structural changes of chromatin on a single cell level remain poorly understood, and it is still largely unclear how the activities of chromatin remodeling complexes are spatially and temporally integrated in living cells. The chromatin ‘opening’ associated with transcriptional activation consists of two distinct changes in chromatin structure: spatial decompaction by changes in nucleosome-nucleosome interactions, and linear decompaction by changes in nucleosome density (Even-Faitelson *et al*., 2016). Both of these processes are thought to be tightly linked to the posttranslational modifications of histones, which characterize transcriptionally active and transcriptionally silent chromatin (Li *et al*., 2007). *In vitro*, specific histone modifications directly influence the spatial organization of chromatin by mediating or restricting nucleosome-nucleosome interactions. For example, the acetylation of histone H4 at lysine 16 prevents the interaction between the acetylated tail and a neighboring nucleosome, lowering the propensity to form a compact 30 nm fiber (Shogren-Knaak *et al*., 2006). However, no effect on linear chromatin compaction was observed on autosomes in mammalian cells and *C. elegans* upon loss of H4K16 modification during differentiation (Taylor *et al*., 2013; Lau *et al*., 2017). In addition, evidence is accumulating that extended regular higher order chromatin structures like a 30 nm fiber do not form *in vivo*, and that chromatin fibers are present mostly in dispersed states in living cells (Fussner *et al*., 2012; Hsieh *et al*., 2015; Chen *et al*., 2016). This is corroborated by recent visualizations of chromatin chains in intact mammalian nuclei, showing that the local appearance of the chromatin fiber is similar in regions of eu- and heterochromatin (Ou *et al*., 2017). It is therefore unclear how local chromatin folding is influenced by histone modifications *in vivo*.

In addition to their potential to mediate inter- and intra-chromosomal interactions and thus to shape the longer-range organization of the chromatin fiber, nucleosomes constitute direct obstacles to the progression of RNA Polymerase II (Pol II) during transcription. Histones and entire nucleosomes are therefore evicted from the DNA during transcription by the concerted action of chromatin remodelers like the SWI/SNF and SWR1 complexes and histone chaperones like Asf1 and FACT (reviewed in (Venkatesh and Workman, 2015)). This nucleosome eviction (leading to a lower nucleosome density) would be expected to cause an increase in the effective length of the chromatin fiber, but this has not been investigated *in vivo*.

In order to study chromatin dynamics and organization *in vivo*, we took advantage of the budding yeast *GAL* locus, a highly regulated gene cluster which has served as a paradigm for inducible gene expression. The *GAL* locus comprises the *GAL7, GAL10* and *GAL1* genes located next to each other on chromosome II. These three genes, encoding enzymes required for the metabolism of galactose, are highly regulated depending on the carbon sources present in the growth medium. The genes are repressed in the presence of glucose, active in the presence of galactose (but absence of glucose) and ‘derepressed’ in the absence of both glucose and galactose (e.g. in raffinose-containing medium). Intricate regulation of carbon metabolic genes allows *Saccharomyces cerevisiae* to adapt to and successfully compete with other organisms for various sugars present in the environment (New *et al*., 2014).

*GAL* gene activation involves the recruitment of several histone modifying enzymes (Carrozza *et al*., 2002; Wang *et al*., 2002; Govind *et al*., 2007) and dramatic reduction of nucleosome occupancy at the locus (Schwabish and Struhl, 2004; Govind *et al*., 2007; Bryant *et al*., 2008). How this affects the chromatin conformation *in vivo* and to which extent chromatin decompaction and transcriptional activation are interdependent remains unclear. To address these important questions, we developed an assay to visualize chromatin compaction in living cells. Monitoring the distance between two chromosomal loci on either side of the *GAL* locus revealed a drastic linear decompaction upon activation of the *GAL* gene cluster. This decompaction was tightly coupled to transcriptional activity. Furthermore, the observed opening was not regulated by histone acetylation but depended on the activity of nucleosome-evicting chromatin remodelers.

## Results

### An assay to quantitatively analyze transcription-induced chromatin decompaction in living cells

To probe chromatin decompaction during transcription in a quantitative manner in living cells, we developed a microscopy-based assay to follow chromatin conformation in S. *cerevisiae* over time. We chose the *GAL7-GAL10-GAL1* gene cluster as a model system, since it is very well studied and the presence of three co-regulated genes spanning ~ 5.8 kb is expected to give a clear decompaction response. LacO and TetO repeats were introduced on either side of the *GAL* gene cluster or in a control region and visualized with LacI-GFP and TetR-mCherry (Figure 1A). Bright green and red dots were readily detected in all cells (Figure 1A) and their positions were determined using fitting of a Gaussian profile to obtain sub-pixel resolution of < 20 nm in x and y and ~ 35 nm in z (Supplemental Figure 1A). Although imaging of live cells introduces potential motion errors, these did not contribute significantly to our measurements (Supplemental Figures 1B and C). The distance between LacI-GFP and TetR-mCherry tagged loci could thus be determined with high precision and was analyzed in cells grown in raffinose (GAL locus derepressed) or stimulated by the addition of glucose (repressed) or galactose (active). In a control region adjacent to but not spanning the *GAL* locus (18 kb between LacO- and TetO-repeat insertion sites), no change in the distribution of measured distances was observed between the different carbon sources (Figure 1B). In contrast, in strains with a distance of 14 or 31 kb between the insertion points of the repeats across the *GAL* locus, the distribution clearly shifted towards larger distances in activated cultures (Figure 1B and C) increasing from a mean distance of 280±9 nm in raffinose to 410±15 nm in galactose in the 14 kb reporter strain (n=7) and from 370±4 nm to 488±8 nm in the 31 kb reporter strain (n=10) (3D distances; values are means of biological replicates with standard error of the mean). As expected, the deletion of the transcriptional activator Gal4 prevented the induction of the *GAL* gene transcription and also abolished the decompaction response, whereas deletion of the transcriptional repressor Gal80 led to transcription and decompaction already in raffinose-grown cells (Supplemental Figure 2A & B). Thus, we have developed a microscopy assay that can readily detect chromatin decompaction upon transcriptional activation in living cells.

**Figure 1:**
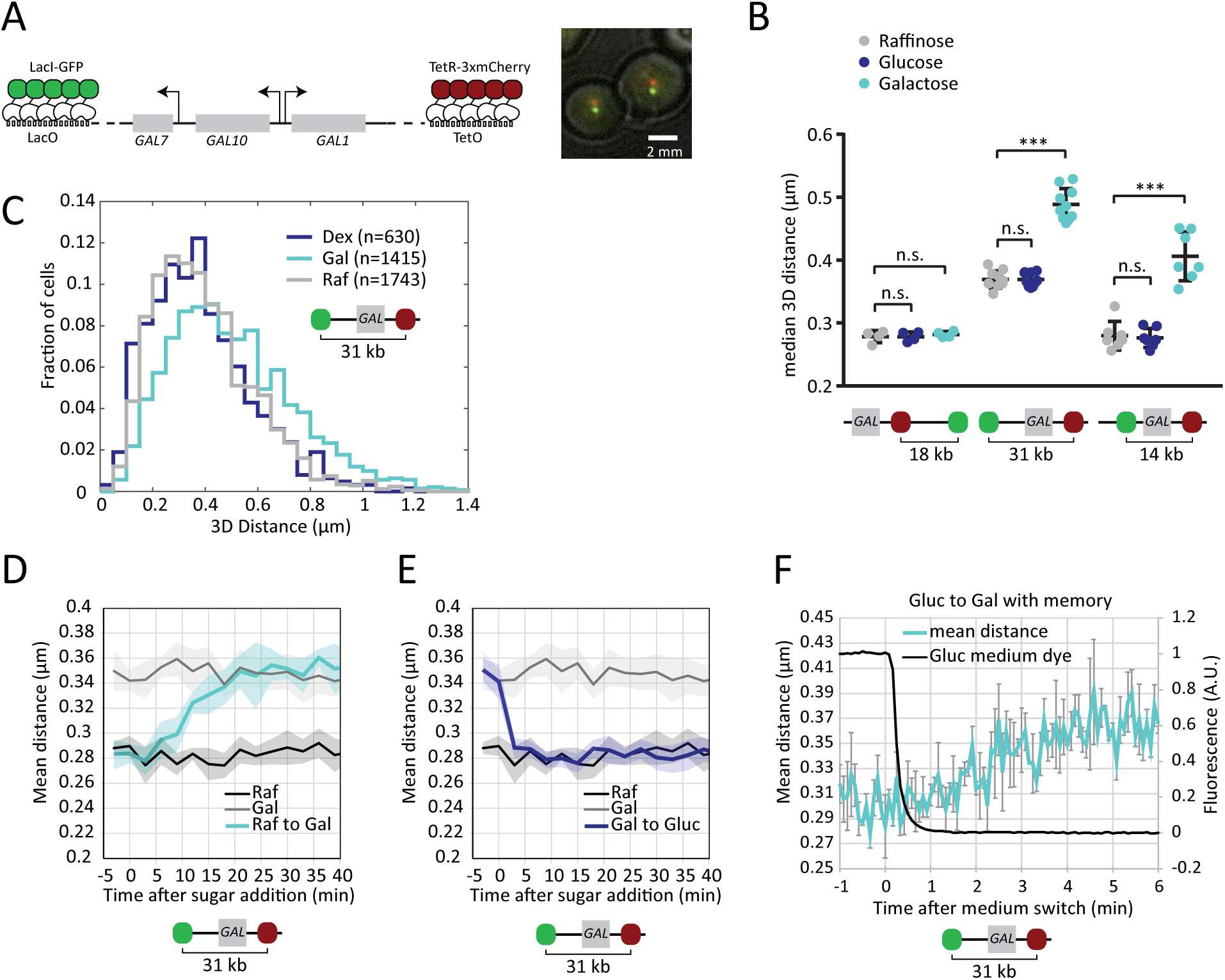
Measuring transcription-dependent chromatin decompaction *in vivo*. (A) Chromosome labeling scheme with LacO and TetO repeats on either side of the *GAL7-10-1* locus and example image of cells expressing the reporter constructs. (B) The distance between LacO and TetO repeats in cell populations one hour after addition of the indicated carbon source to the raffinose-containing culture medium. Each dot represents the median from an independent experiment with at least 100 analyzed cells (typically 600-1000). Lines and error bars indicate means and standard deviations. *** adjusted p-value < 0.001, n.s. not significant (p>0.05). (C) Histogram of distance distribution for TetR and LacI dots for 31 kb reporter strain in raffinose (grey), glucose (dark blue) or galactose (light blue). Shown is one biological replicate from Figure 1B. (D) Mean distance of cell population during addition of galactose to cells grown in raffinose (or continuous growth in either galactose or raffinose). Shown is the mean of four independent experiments, shaded areas indicate the standard error (SE). In each experiment > 100 cells were analyzed per timepoint. (E) Mean distance of cell population during addition of glucose to cells grown in galactose (or continuous growth in either galactose or raffinose). Shown is the mean of four independent experiments, shaded areas indicate the standard error (SE). In each experiment > 100 cells were analyzed per timepoint. (F) Mean distance of cell population during medium switch from glucose to galactose. Cells were pre-induced by overnight growth in galactose and then grown in glucose for one hour. Black curve shows fluorescence of Dextran-Alexa Fluor 680 (3000 MW, anionic) present in the glucose medium to monitor the medium switch in the microfluidics setup. Shown is the mean of three independent experiments, error bars indicate the standard error (SE). Note high fluctuation due to low cell numbers (30-50 per timepoint and experiment).

Next, we analyzed the dynamics of decompaction in time-lapse experiments. For time-lapse experiments, we only acquired single plane images and analyzed 2D distances. This simplified assay allowed us to process all control strains and conditions in a single experiment yet still provided sufficient sensitivity to detect changes in distance as observed in 3D (Figure 1D-F). Acute induction of the *GAL* genes by addition of galactose to cells growing in raffinose led to an increase in the population mean distance over the course of 20 minutes (Figure 1D) mimicking the kinetics of transcriptional activation as seen e.g. by quantitative PCR (qPCR) analysis (Green *et al*., 2012). Similarly, the addition of glucose to cells growing in galactose resulted in fast compaction (Figure 1E) corresponding to the rapid shut-down of transcription due to glucose-induced repression. Decompaction could also be induced with faster kinetic within less than five minutes if cells were pre-grown in galactose and then repressed with glucose for only 1 h before re-induction with galactose (Figure 1F). In this case, the presence of elevated levels of Gal1 and Gal3 proteins induce a ‘transcriptional memory’ and lead to faster activation (Kundu and Peterson, 2010). Thus, the kinetics of chromatin de- and re-compaction closely follow transcriptional activity.

### Active transcription correlates with chromatin decompaction in single cells

To directly correlate transcriptional activity with distance measurements on a single cell level, we applied single molecule fluorescent *in situ* hybridization (smFISH) to visualize transcripts as they are produced from the *GAL* locus in our distance reporter strains. We simultaneously used FISH probes for all three *GAL* genes *GAL1, GAL10* and *GAL7*, all labeled with the same fluorophore (Qasar670). While cytoplasmic mRNAs are visible as individual spots or as hazy signal due to the large number of molecules in induced cells, the transcription site is visible as a bright focus in the nucleus close to the repeat-marked gene loci (Figure 2A). Transcription spots and cytoplasmic mRNA foci are absent in cells grown in raffinose (Figure 2A, first panel). Upon induction with galactose, the fraction of cells with transcription spots increased over the course of 30 min mirroring the increase in the mean distance in the population after induction (Figure 2B). After 60 minutes or longer, a transcription spot can be seen in virtually every cell (Figure 2A), but since our automatic detection routine does not detect all transcription spots, the maximum measured percentage of cells with transcription spot was only ~ 80 % (Figure 2B).

**Figure 2:**
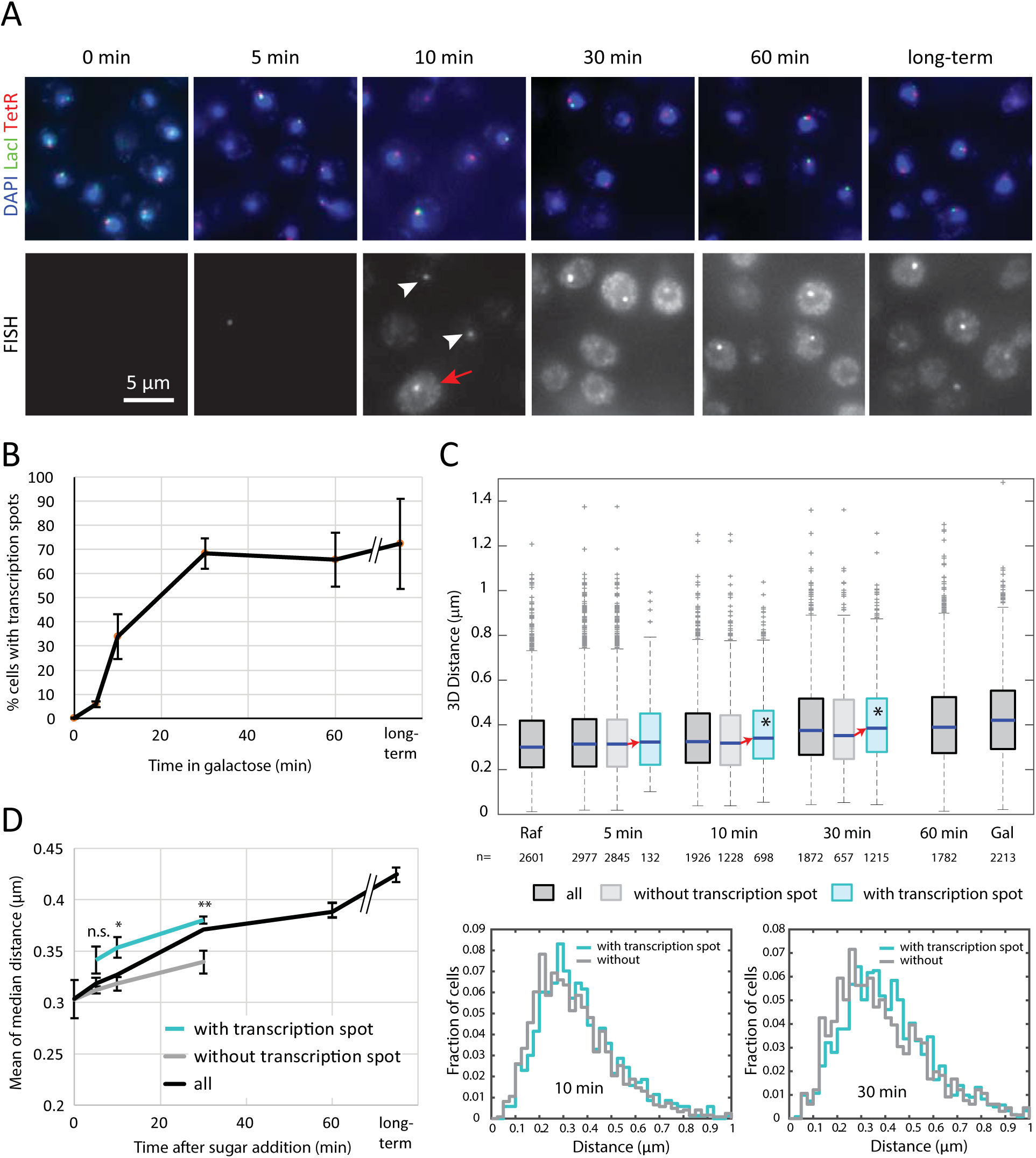
Chromatin opening correlates with transcription on single cell level. (A) Individual slices of image stacks showing single molecule FISH signal (bottom) in cells expressing genomic markers (green and red) and stained with DAPI (blue) grown in raffinose (0 min) or after different times of induction with galactose. The cells were hybridized with probes against *GAL1, GAL7* and *GAL10* mRNAs simultaneously, all labeled with the far-red dye Qasar670. In the image of the 10 min time point, two transcription spots in cells with little cytoplasmic mRNA signal are indicated by white arrowheads, and a cell with many cytoplasmic mRNA foci is indicated with a red arrow. (B) Quantification of percentage of cells in which a bright transcription spot could be detected (Mean of three biological replicates, error bars indicate standard deviation). (C) Distribution of distances across the *GAL* locus for all cells (dark grey) and cells without (light grey) or with (light blue) transcription spot. Stars indicate that median of the distances in cells classified as carrying a transcription spot is outside the 95 % confidence interval of the median of the distances in cells classified as not carrying a transcription spot and vice versa. n: number of cells analyzed. Histograms below show comparison of distance distributions at 10 min and 30 min after induction. Shown is the data of one representative experiment out of three biological replicates. (D) Means of median distances in cells classified for absence or presence of a transcription spot from three biological replicates (Error bars represent standard deviation). Difference between population with and without spots was tested using an unpaired 2-tailed t-test not assuming equal variance. n.s. p>0.05, * p<0.05, **p<0.01.

To correlate transcription with distance, we analyzed early time points after induction, when only a fraction of the cells showed a transcription spot (Figure 2B). We separately determined the 3D distance between the LacO- and TetO-repeat markers in cells with or without a detectable transcription spot. 5 min after induction, decompaction was still minimal and we observed only few cells with a transcription spot. However, after 10 and 30 min the distances in cells with a transcription spots were significantly larger than in cells without a transcription spot (Figure 2C and D). Thus, decompaction correlates with transcription also on a single cell level.

After transcriptional repression by the addition of glucose, the signal of *GAL* mRNAs in the cytoplasm quickly diminished (Supplemental Figure 3A). Although the nuclear FISH signal at the site of transcription was reduced as well, a weak spot could still be detected in many cells even 60 minutes after glucose addition (Supplemental Figure 3A and C). Most likely this corresponds to poly-adenylated *GAL* mRNAs that were previously reported to persist at the *GAL* gene loci upon transcriptional shut-off (Abruzzi *et al*., 2006). Interestingly, cells with a brighter RNA dots showed a distance distribution shifted to longer distances compared to cells with a weaker or absent RNA dot signal (Supplemental Figure 3B). This might indicate that transcription is still ongoing in those cells and thus the brighter RNA signal could stem from nascent transcripts. Alternatively, long-lived, chromatin-associated RNAs might contribute to keeping chromatin in a partially decompacted state.

### Histone acetylation is dispensable for chromatin decompaction at the *GAL* locus

The increased distance distribution observed in our assay, which is indicative of decompaction of chromatin during transcription, could originate from reduced internucleosomal interactions, from the eviction of nucleosomes, or from both. Internucleosomal interactions are often mediated by histone tails and are thought to be regulated by posttranslational modification. Histones H3 and H4 contribute most to direct internucleosomal interactions but are also required for the recruitment of transcriptional activators. Acetylation of H3 *in vivo* occurs predominantly through Gcn5 (Grant *et al*., 1997). This histone acetyl transferase is a component of the SAGA complex, which is recruited to the *GAL* locus by the transcriptional activator Gal4 (Carrozza *et al*., 2002; Govind *et al*., 2007). However, neither the deletion of the SAGA histone acetyl transferase Gcn5 nor that of its activating subunit Ada2 influenced chromatin decompaction across the *GAL* locus (Figure 3A). These mutations also do not affect pre-initiation complex formation at the *GAL* promoter (Bhaumik and Green, 2001) or mRNA production (Stafford and Morse, 2001; Green *et al*., 2012; Figure 3B), suggesting that acetylation is dispensable for transcriptional activation and chromatin decompaction induced by Gal4 (Note that mRNA expression data normalized to the same conditions in the wild type strain rather than to the raffinose condition in the same strain are shown in Supplemental Figure 4.). While dispensable for activation, Gcn5 activity is crucial for the association of the *GAL* locus with the nuclear periphery upon activation (Dultz *et al*., 2016). Therefore, our results rule out that the observed decompaction response is a consequence of relocalization of the *GAL* locus to the nuclear pore complex, which might, for example, generate pulling forces that could lead to stretching of the chromatin fiber.

**Figure 3:**
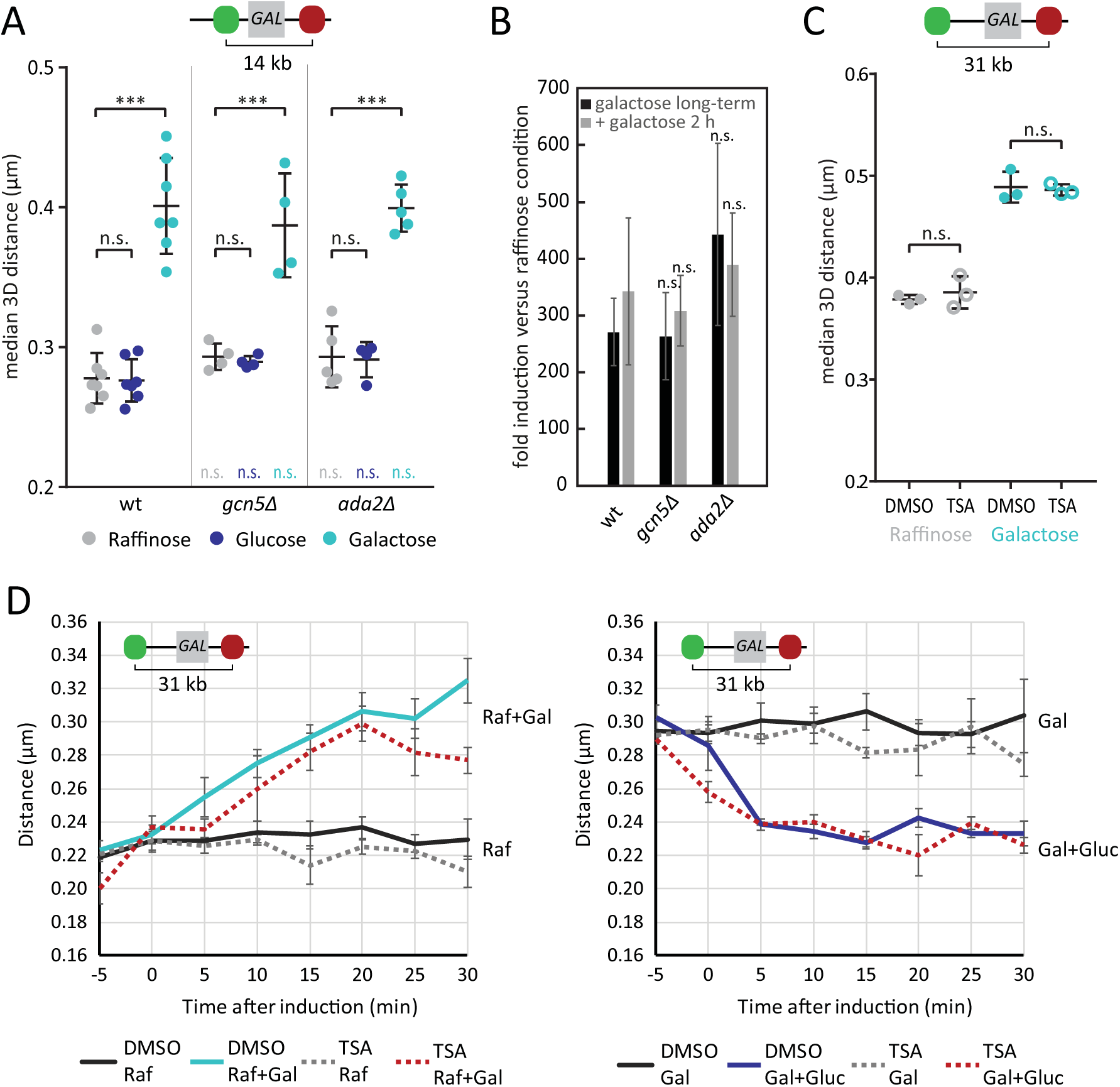
The role of acetylation in chromatin opening. (A) The median distance between LacO and TetO repeats in cell populations one hour after addition of the indicated carbon source to the raffinose containing culture medium to a final concentration of 2% (raffinose 4%). Each dot represents the median of an independent experiment with at least 200 analyzed cells (typically 600-1000). Lines and error bars indicate means and standard deviations. *** adjusted p-value < 0.001,n.s. not significant (p>0.05). Colored n.s. indicate that there is not significant difference to the wild type (wt) in the same condition. (B) Fold induction of *GAL10* mRNA compared to growth in raffinose determined by quantitative PCR (normalized for *ACT1* mRNA as endogenous control). Means of 3-4 biological replicates are shown. Error bars represent SE. (C) The median distance between LacO and TetO repeats in cell populations at steady state growth in medium containing 2% galactose or 2% raffinose. Each dot represents the median of an independent experiment with at least 300 analyzed cells (typically 600-1000). Lines and error bars indicate means and standard deviations. n.s. not significant. (D) Opening and closing kinetics in the presence of TSA or DMSO. Cultures grown for 16 hours in the presence of DMSO or TSA were diluted 1:1 with medium containing TSA or DMSO respectively as well as 4% glucose or galactose to obtain a final concentration of 2% in the medium. Mean of three biological replicates is shown, error bars represent standard error of the mean. > 100 cells were analyzed per timepoint in each condition and experiment.

We also tested whether the inhibition of histone deacetylases using trichostatin A (TSA) would affect chromatin decompaction. TSA treatment neither changed the steady state distance distributions (Figure 3C) nor did it affect induction or repression kinetics (Figure 3D). Since both the deletion of *GCN5* and the inhibition of histone deacetylases targeted by TSA globally affect acetylation levels of histones (Vogelauer *et al*., 2000 and Supplemental Figure 5A), these results indicate that the global acetylation state of histones *per se* does not play a major role in regulating chromatin compaction at the *GAL* locus as detected by our assay.

In contrast to the *GAL* genes, other budding yeast genes were previously shown to fully depend on the HAT activity of Gcn5 for their transcriptional activation including a β-estradiol responsive transcriptional activator based on the viral VP16 transactivation domain (Stafford and Morse, 2001). Intriguingly, we found that upon activation via a β-estradiol-inducible VP16 fusion protein (Louvion *et al*., 1993), Gcn5 was essential for decompaction (Supplemental Figure 5B-D) and neither *gcn5Δ* nor *spt20Δ* cells displayed decompaction of the *GAL* locus upon addition of β-estradiol (Supplemental Figure 5E&F), suggesting that transcriptional activation and decompaction are closely linked. These data also show that the observed decompaction response is not specific for Gal4-mediated activation.

### Mutants that affect transcription also affect decompaction

Although Gcn5 was dispensable for chromatin decompaction, deletion of the SAGA component Spt20, which is required for the stability of the SAGA complex (Grant *et al*., 1997) and for its recruitment by Gal4 (Bhaumik and Green, 2001; Larschan and Winston, 2001; Bryant and Ptashne, 2003), led to strongly reduced decompaction of the *GAL* locus (Figure 4 A and C). As previously reported (Bhaumik and Green, 2001), Spt20 mutants also show strongly reduced *GAL* gene transcription (Figure 4B).

**Figure 4:**
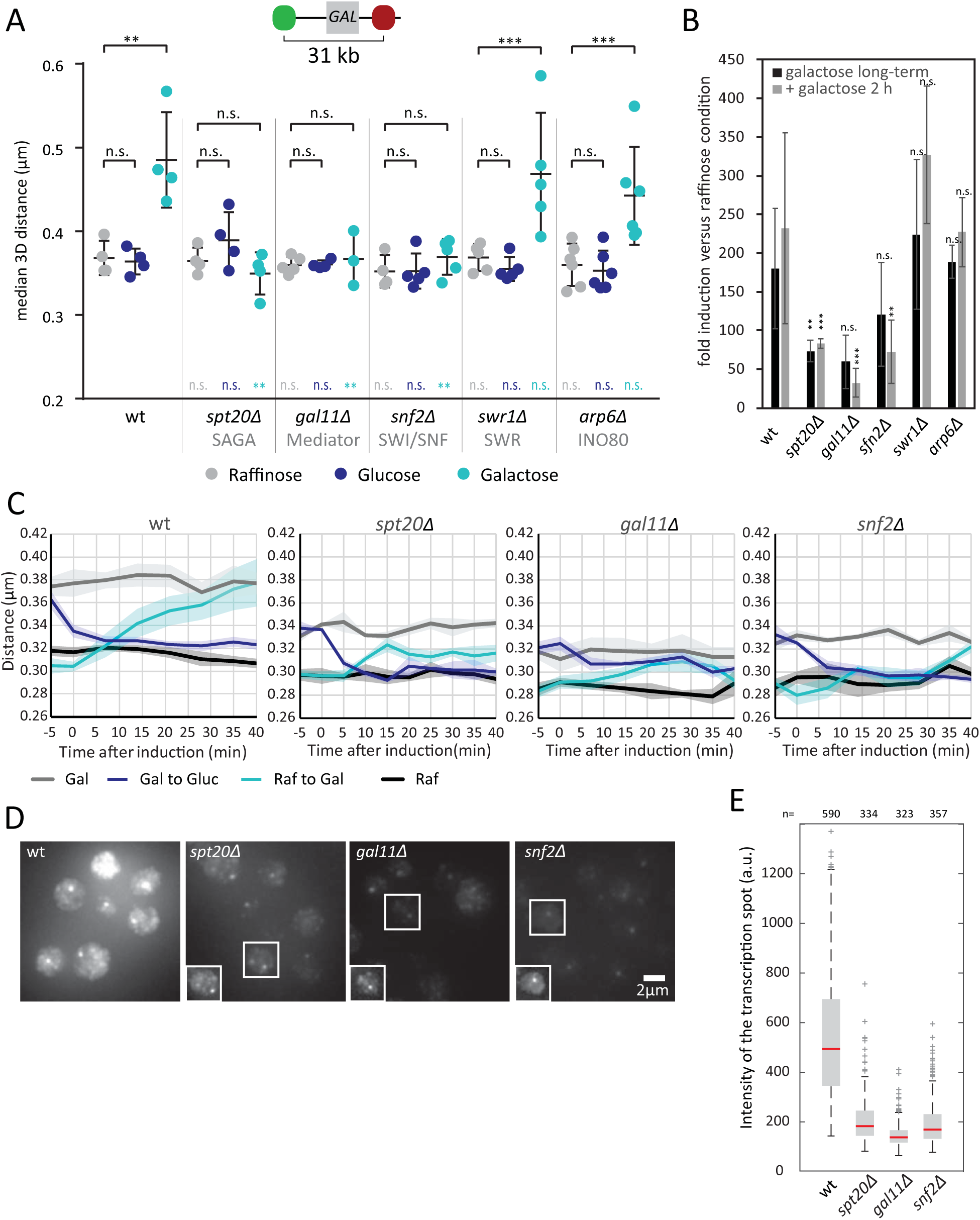
Decompaction correlates with transcriptional activation. (A) Distance between LacO and TetO repeats in cell populations one hour after addition of the indicated carbon source to the raffinose containing culture medium to a final concentration of 2% (raffinose 4%). Each dot represents the median of an independent experiment with at least 100 analyzed cells (typically 600-1000). Lines and error bars indicate means and standard deviations. ** adjusted p-value < 0.01, n.s. not significant (p > 0.05). Colored p-values indicate significance relative to the wild type (wt) in the same condition. (B) Fold induction of *GAL10* mRNA compared to growth in raffinose determined by quantitative PCR (normalized for *ACT1* mRNA as endogenous control). Means of three biological replicates are shown. Error bars represent SE. (C) Kinetics of chromatin opening and closing in wild type and mutant cells. Cells were grown overnight in medium with 2 % raffinose or 2 % galactose. Galactose or glucose respectively were added or not added to 2 % final concentration at time point 0. Shown are the means of three or more biological replicates with the standard error represented by the shaded areas. (D) smFISH for *GAL1, GAL7* and *GAL10* in the indicated strains growing at steady-state in galactose. Insets show enhanced contrast for indicated cells. Images are maximum intensity projections of z stacks encompassing the entire cells. (E) Boxplot showing the intensities of transcriptions spots in the different strains (distribution from one representative experiment out of two biological replicates).

Whereas SAGA is thought to be required to stabilize the general transcriptional coactivator Mediator at the *GAL* promoter, Gal4 is also able to directly recruit this complex (Traven *et al*., 2006). Indeed, cells with a deletion of *GAL11*, a Mediator component that mediates direct interaction with Gal4, or expressing a temperature sensitive mutant of the Mediator protein Med7 *(med7-163)* phenocopied *spt20A* cells by exhibiting reduced overall decompaction in galactose as well as slowed decompaction kinetics (Figure 4 A and C and Supplemental Figure 6).

To analyze the effect of *SPT20* or *GAL11* deletions on a single cell level, we carried out single molecule FISH. After overnight growth in galactose, most cells displayed cytoplasmic mRNA signals, but the signal was strongly reduced compared to wild type cells (Figure 4D). In addition, the intensity of transcription spots, which were still present in most cells, were reduced on average to ~ 30 % of the wild type intensity (Figure 4D and E). Thus, also in the mutant cells, reduced transcription correlates with reduced decompaction of the *GAL* locus. Our results reveal a close linkage between transcriptional activation and decompaction, but they are not sufficient to establish a causality or temporal order between the two processes.

We also attempted to directly address the role of Pol II in decompaction using cells expressing a temperature-sensitive mutant allele of the largest subunit of RNA Pol II, *rpb1-1. rpb1-1* cells were induced with galactose one hour after a shift to the restrictive temperature. Both expression of *GAL10* mRNA and decompaction were reduced but not abolished (Supplemental Figure 7A and B). Thus, it appears that transcriptional induction at the *GAL* locus is so strong that the temperature-sensitive *rpb1-1* mutant is not fully effective in suppressing decompaction and transcription.

### Nucleosome remodelers are required for decompaction

Next, we examined the function of nucleosome remodelers in our decompaction assay. As demonstrated by chromatin immunoprecipitation experiments, histone occupancy in the promoter of transcribed genes – but also in the open reading frame (ORF) of highly transcribed Pol II genes including the *GAL* genes – is reduced several fold upon activation (Lee *et al*., 2004; Schwabish and Struhl, 2004) (Xin *et al*., 2009). Nucleosome eviction is mediated by the SWI/SNF chromatin remodeling complex and the histone chaperones Asf1 (Antisilencing function 1) and FACT (FAcilitates Chromatin Transcription), a heterodimeric complex consisting of Pob3 and Spt16 (Schwabish and Struhl, 2006, 2007; Venkatesh and Workman, 2015). If the observed decompaction across the *GAL* locus is due to nucleosome eviction, depletion of either of these factors should lead to a reduction of decompaction.

SWI/SNF is a chromatin remodeling complex with nucleosome sliding and nucleosome eviction activities. At the *GAL* locus, it is required for rapid Pol II recruitment and activation (Kundu and Peterson, 2010). We found that deletion of *SNF2*, encoding the ATPase subunit of the complex, slowed down the kinetics of decompaction and strongly reduced the degree of final decompaction as well (Figure 4A and C) consistent with a reduction in nucleosome eviction across the locus (Schwabish and Struhl, 2006). Although inactivation of SWI/SNF has previously been reported not to alter steady-state *GAL* gene expression (Lemieux and Gaudreau, 2004), we observed reduced mRNA levels both by qPCR and smFISH in *snf2Δ* cells (Figure 4B and D). Again, transcriptional activation and decompaction correlated in this mutant, and we observed both reduced transcription and reduced decompaction. In contrast to the SWI/SNF complex, the chromatin remodeling complexes INO80 (represented by arp6Δ) and SWR1 (represented by swr1Δ) were neither required for chromatin decompaction nor transcriptional activation at the *GAL* locus (Figure 4A and B).

Next, we tested the involvement of the histone chaperones Asf1 and FACT, which have both been implicated in the regulation of the *GAL* genes (Schwabish and Struhl, 2006; Xin *et al*., 2009). Deletion of *ASF1* did not prevent chromatin decompaction in our assay (Figure 5A), which is consistent with previous findings that showed only a small decrease in histone H3 (but not H2B) removal at *GAL* promoters and ORFs (Schwabish and Struhl, 2006). Deletion of ASF1 also did not affect transcriptional activation of *GAL10* (Figure 5B). In contrast, cells carrying temperature sensitive alleles for either component of the FACT complex (pob3-7 or spt16ts) showed defects in decompaction one hour after a shift to activating conditions at the restrictive temperature (Figure 5D). While pob3-7 cells exhibited slightly increased distances over the GAL locus already in raffinose and only a moderate additional increase upon activation with galactose, *spt16ts* cells showed no decompaction at the restrictive temperature. Importantly, both strains showed similar levels of GAL10 mRNA induction as wild type cells at both permissive and restrictive temperatures (Figure 5C. Note that both mutant and wild type cells exhibited reduced *GAL10* mRNA induction at 37 °C compared to 25 °C). Therefore, in the *spt16ts* strain, decompaction and transcription are uncoupled, showing that transcription can occur in the absence of detectable decompaction. Although transcription at 37 °C might not proceed as efficiently as at 25 °C, there is no added defect in FACT mutants which are clearly defective in decompaction. Together, these findings indicate that the activities of SWI/SNF and FACT are major sources of decompaction and suggest that nucleosome eviction leads to the increase in distance that we observe across the activated locus. However, we cannot exclude that processes other than nucleosome eviction and associated with actively transcribing Pol II including histone tail modifications contribute to the observed decompaction as well. Importantly, the *spt16ts* mutant shows that the observed decompaction is not required for at least moderate levels of transcription.

**Figure 5:**
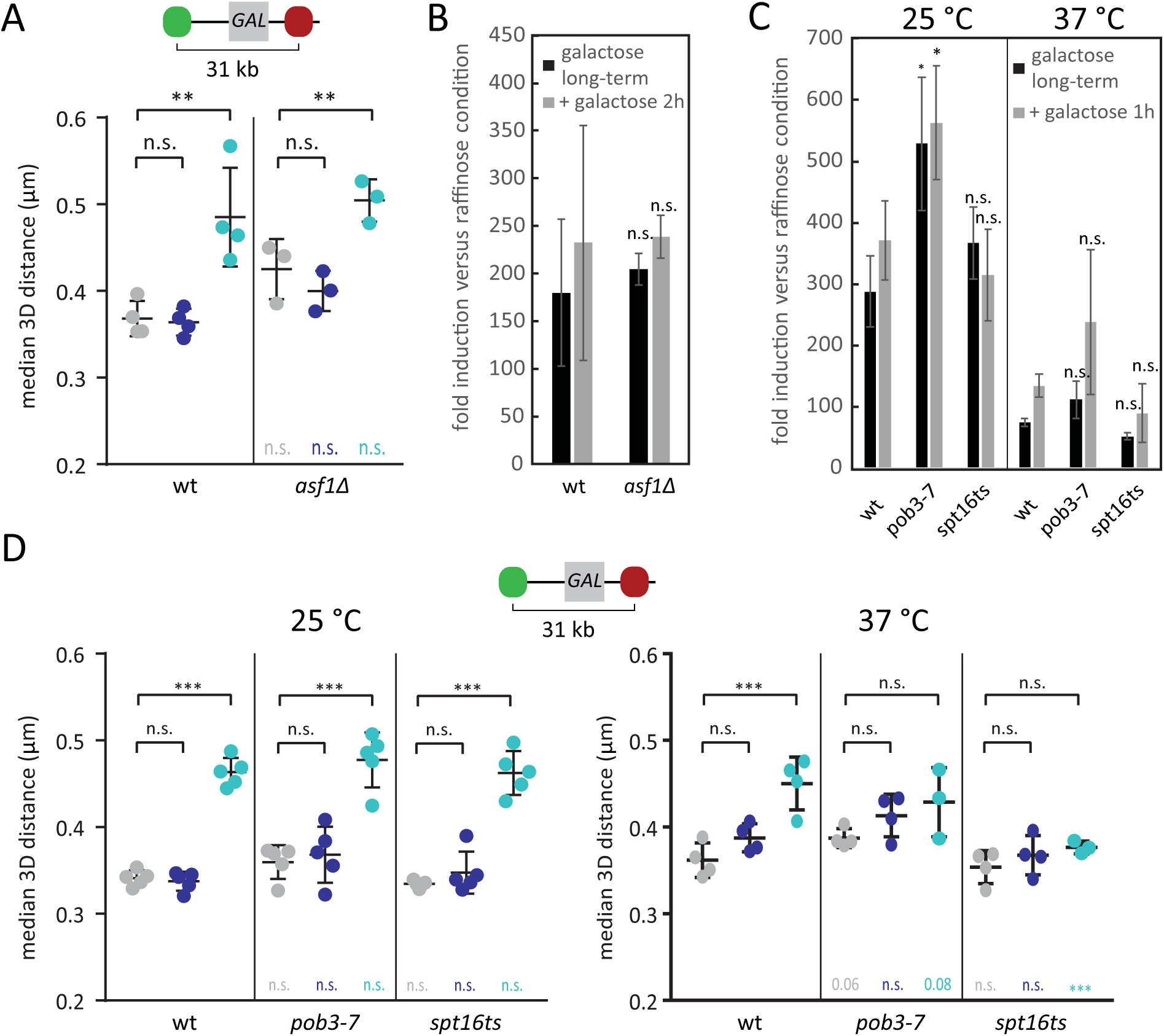
Nucleosome chaperones are crucial for chromatin opening. (A) Distance between LacO and TetO repeats in cell populations one hour after addition of the indicated carbon source to the raffinose containing culture medium to a final concentration of 2% (raffinose 4%). Each dot represents the median of an independent experiment with at least 100 analyzed cells (typically 600-1000). Lines and error bars indicate means and standard deviations. *** adjusted p-value < 0.001, n.s. not significant (p > 0.05). Colored p-values indicate significance relative to the wild type (wt) in the same condition. (B) Fold induction of *GAL10* mRNA in wild type and *asf1Δ* cells relative to raffinose growth conditions determined by quantitative RT-PCR (normalized *for ACT1* mRNA as endogenous control). Means of 3-4 biological replicates are shown. Error bars represent standard errors. (C) Fold induction of *GAL10* mRNA in wild type cells and cells carrying temperature-sensitive allels of FACT components relative to raffinose growth conditions determined by quantitative RT-PCR (normalized for *ACT1* mRNA as endogenous control). Means of 3-4 biological replicates are shown. Error bars represent standard errors. (D) Distance between LacO and TetO repeats in cell populations one hour after addition of the indicated carbon source to the raffinose containing culture medium to a final concentration of 2% (raffinose 4%). Cells were cultured in raffinose medium at 25 °C and shifted to 37 °C for the indicated experiments one hour prior to addition of the different carbon sources. Each dot represents the median of an independent experiment with at least 200 analyzed cells (typically 600-1000). Lines and error bars indicate means and standard deviations. ** adjusted p-value < 0.01, n.s. not significant (p > 0.05). Colored p-values indicate significance relative to the wild type (wt) in the same condition.

### Decompaction of the *GAL* locus is transcription dependent

If the observed decompaction is indeed due to the removal of nucleosomes at the promoter and the ORF, the degree of decompaction is expected to scale with the length of the ORF. To test this prediction, we introduced *GAL* promoter driven reporter genes at a different genomic site ~ 10 kb telomeric of the native *GAL* locus and measured changes in chromatin distance in the presence or absence of galactose (Figure 6). The *GAL* promoter immediately followed by a terminator sequence did not exhibit a significant decompaction response. In contrast, robust decompaction was observed in a strain where the entire *GAL1* gene including promoter and downstream sequences was inserted. Decompaction, although to a weaker extent than for the entire *GAL1* gene, was also observed in a strain where we introduced a *GAL1* promoter-driven ORF coding for Glutathione S transferase (GST). Since the ORF of a single *GST* is shorter than the one of *GAL1* (700 bp versus 1 kb), we also examined the response in a reporter strain carrying two consecutive copies of the *GST* ORF *(2xGST)*. The presence of a second *GST* ORF increased the decompaction to a similar extent as the *GAL1* ORF suggesting that decompaction does indeed scale with ORF length, although a contribution of internal or 3’UTR sequences cannot be excluded.

**Figure 6:**
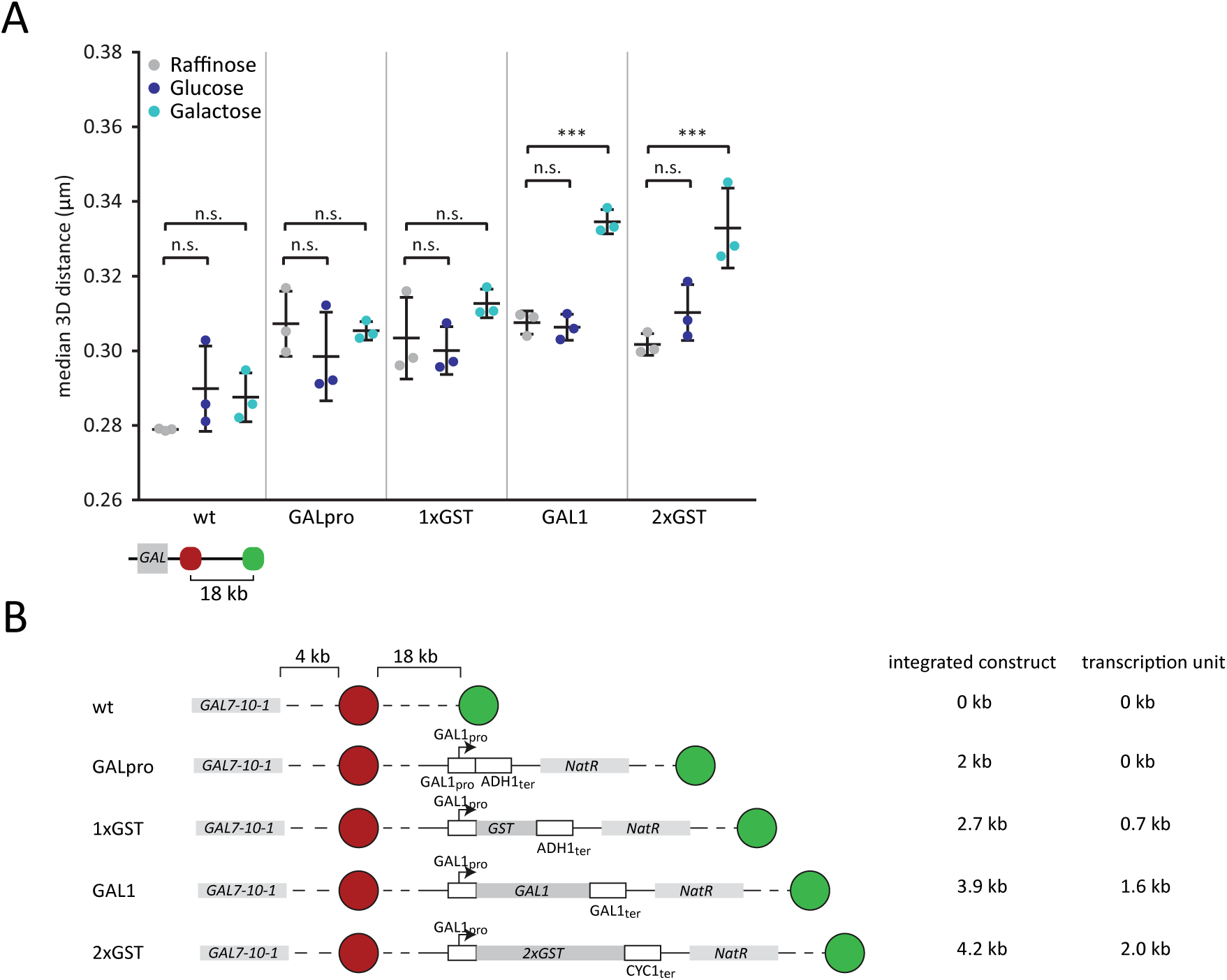
Dependence of 3D distance on ORF length. (A) The median distance between LacO and TetO repeats in cell populations one hour after addition of the indicated carbon source to the raffinose containing culture medium. Each dot represents the median from an independent experiment with at least 100 analyzed cells (typically 600-1000). Lines and error bars indicate means and standard deviations. *** adjusted p-value < 0.001, n.s. not significant (p > 0.05). (B) Schematic representation of constructs integrated. Wild type strain (wt) corresponds to control strain shown in Figure 1B.

### Decompaction leads to an untypical open chromatin state at the *GAL* locus

Our data provide the first quantitative measurement of transcription-dependent decompaction in living cells. To understand the changes in chromatin structure underlying the observed decompaction response, we analyzed the distance distributions of the populations more closely. The 3D distance of genomic loci on the same chromosome has been shown to scale with the distance in base pairs (van den Engh *et al*., 1992; Arbona *et al*., 2017). This is also true on chromosome II of budding yeast in both glucose and galactose grown cells as shown previously by us (Dultz *et al*., 2016 and Figure 7A grey and black data points). One explanation for the observed changes in compaction at the *GAL* locus could be that the *GAL* locus – compared to bulk chromatin – is hyper-compacted in the repressed state. However, this is not the case: in the repressed or de-repressed state (glucose or raffinose growth), the median distances of TetO and LacO marker pairs across the *GAL* locus (Figure 7A open red circles, data for raffinose grown cells not shown) are in line with the distances of other pairs of loci on chromosome II. This indicates that the compaction state of the locus is comparable to other regions on the same chromosome. In contrast, in galactose-grown cells, the median distance increased far above the distance expected from the linear distance on the chromosome (Figure 7A, filled red circles), showing that the *GAL* locus is strongly hypo-compacted in its active state. This is consistent with the interpretation that linear decompaction at the *GAL* locus upon transcriptional activation is due to the eviction of nucleosomes, and that the high transcriptional activity over three clustered genes at the *GAL* locus leads to a very low histone density and thus hypo-compaction.

**Figure 7:**
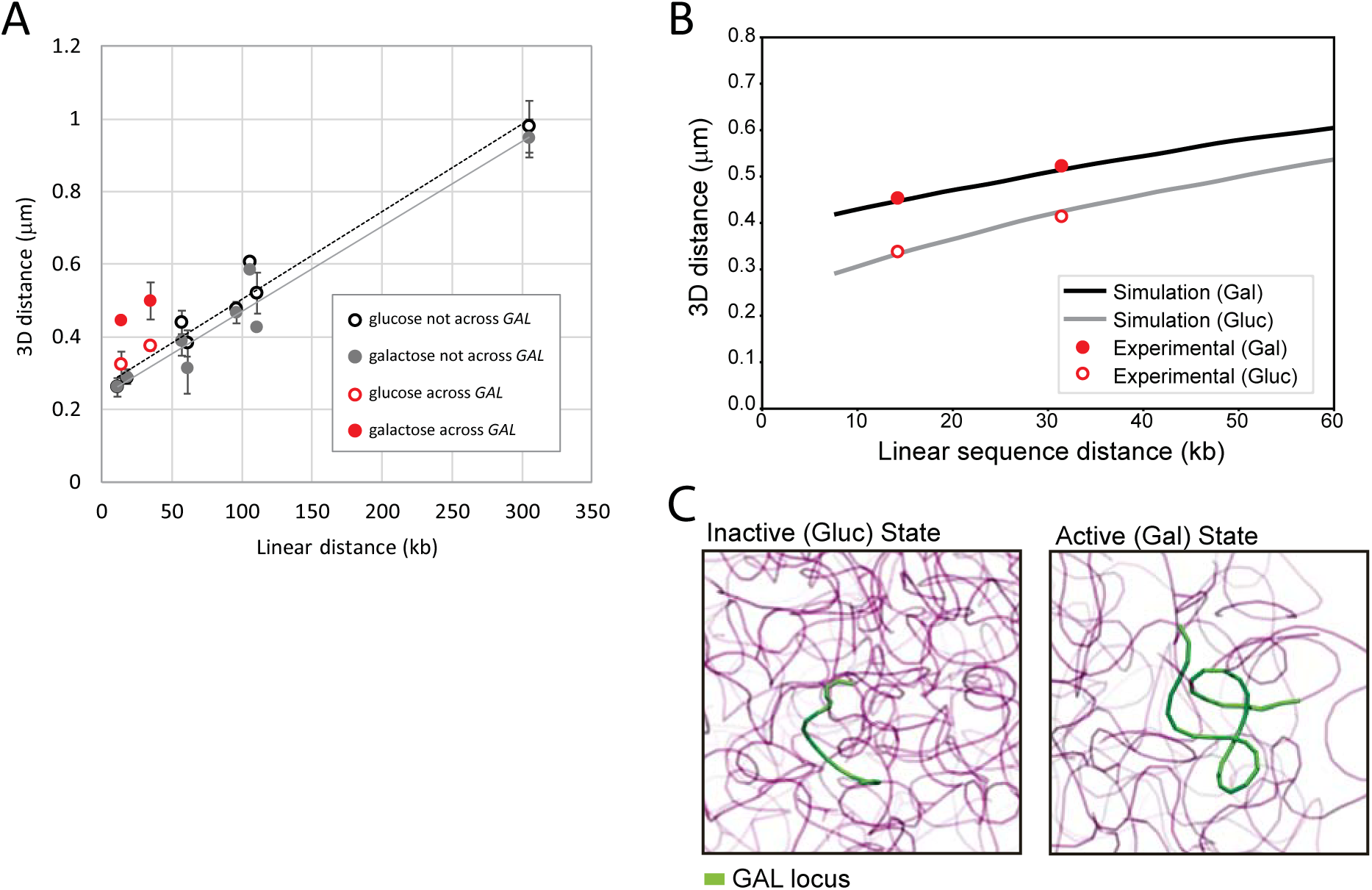
Dependence of 3D distance on the linear distance along the chromosome. (A) Experimentally determined mean 3D distance in various strains carrying the chromatin location markers TetO and LacO at different distances from each other on chromosome II. Each datapoint is the mean of three or more medians from different biological replicates with the standard deviation indicated by the error bars. Grey and black datapoints are plotted from data published in Dultz *et al*., 2016 Figure 4B. (B) Average 3D distance vs. genomic distance curves obtained from simulated structures at two different values of compaction for the *GAL* locus. The markers correspond to the experimental averages in the active and inactive conditions. In order to reproduce the experimentally observed 3D distances, the compaction of *GAL* locus was set to 200 and 375 bp/bead respectively in the active and inactive conditions. (C) Two randomly selected snapshots of the simulated GAL locus in the inactive (Gluc, left panel) and active (Gal, right panel) conditions.

Since our data provide quantitative measurements of this hypo-compaction, we applied computational modeling to predict the level of histone loss that could lead to such a distance increase. To this end, we adapted a previously developed polymer model of the yeast genome, which accurately recapitulates several experimentally determined features (Tjong *et al*., 2012; Dultz *et al*., 2016), by increasing the resolution of the model in the proximity of the *GAL* locus. In this model, the compaction of the chromatin fiber in the *GAL* locus had to be reduced by ~60% in order to recapitulate the observed increase in distances (Figure 7B and C), which corresponds to a 1.7-fold increase in linear extension. Since the wrapping of DNA on nucleosomes leads to an approximately 6-fold compaction, this suggests that about 30 % of the nucleosomes at the *GAL* locus are evicted upon transcriptional activation. This is in accordance with nucleosome occupancy measurements (Schwabish and Struhl, 2006) and, at a density of 162bp/nucleosome (Horz and Zachau, 1980), corresponds to the eviction of approximately 11 of 36 nucleosomes across the locus.

## Discussion

The recent development of chromosome conformation capture techniques has revolutionized the characterization of three-dimensional interaction landscapes of chromatin and has led to many important discoveries and new models of chromatin organization (reviewed in Pombo and Dillon, 2015; Bonev and Cavalli, 2016). However, these techniques require the fixation of cells and are usually applied to cell populations. Furthermore, it is not possible to directly convert the interactions detected by chromosome conformation capture into 3D distances in the cell nucleus. In fact, a direct comparison of chromosome capture analyses with DNA fluorescence *in situ* hybridisation results has revealed that the two techniques do not always result in congruent findings (Williamson *et al*., 2014). It is therefore crucial to analyze chromatin conformation *in situ* and also *in vivo*.

Chromatin decompaction upon transcriptional activation has been visualized in mammalian cells using large repeat arrays (Tumbar *et al*., 1999; Dietzel *et al*., 2004; Verschure *et al*., 2005; Hu *et al*., 2009). FISH analysis has also been employed to obtain distance measurements at activated loci (Chambeyron and Bickmore, 2004; Chambeyron *et al*., 2005; Morey *et al*., 2007). Although making important contributions to our understanding of changes in chromatin conformation upon transcriptional activation, quantitative interpretation is limited because fixation and DNA FISH protocols introduce modifications to the cell ultrastructure and repeat arrays may themselves influence the local chromatin environment. Therefore, we have used the budding yeast *Saccharomyces cerevisiae* to generate a system where live cell analysis is possible in a quantitative manner at an endogenous gene locus. Although we initially use a 3-gene system to establish our assay, we also show that this assay is sensitive enough to visualize decompaction of a single activated gene. Furthermore, the use of budding yeast allows us to study the basic level of chromatin decompaction in the absence of confounding changes in chromatin conformation like enhancer-promoter looping or looping out from a chromosome territory. Thus, our data yield quantitative measurements of chromatin decompaction upon transcriptional activation and allow us to model the changes in chromatin properties at an activated locus.

### The role of acetylation

Interestingly, we found that deletion of the lysine acetyl transferase gene *GCN5* or inhibition of lysine deacetylation by trichostatin A had no observable effect on the distance distribution in our compaction assay. In contrast, the treatment of mammalian cells with TSA was previously reported to affect chromatin organization globally and to lead to decompaction of heterochromatic regions (Toth *et al*., 2004; Lleres *et al*., 2009). However, these studies used global histone signals (intensity or FRET-FLIM) to assess compaction levels. These analyses thus do not give sequence-specific readout and report on spatial rather than on linear compaction. In agreement with our findings, DNA FISH analysis in mouse cells, which also tested linear compaction between two loci, showed decompaction in polycomb silenced regions but not elsewhere (Eskeland *et al*., 2010). Thus, very low acetylation levels may be crucial for the formation of certain subnuclear compartments in heterochromatic regions of the nucleus but may play a less prominent role in linear decompaction in the relatively open bulk yeast chromatin. Nevertheless, it cannot be excluded that specific acetylation marks added by acetyl transferases other than Gcn5 or removed by non-TSA sensitive deacetylases contributes to linear chromatin decompaction. Furthermore, acetylation could facilitate the binding of factors to specific DNA sequences by rendering the interactions between histones and DNA less tight. This could allow nucleosomes to slide more easily or create binding sites for bromodomain-containing chromatin effector proteins (Filippakopoulos and Knapp, 2014).

### The interplay of transcription and decompaction

Our mutant analysis shows that transcriptional activation and chromatin decompaction are tightly coupled. Unfortunately, we could not directly analyze the requirement of transcription *per se*, since the *rpb1-1* mutant surprisingly did not fully suppress *GAL* gene activation (and decompaction; Supplemental Figure 7). Nevertheless, together with the observation that an ectopic *GAL* promoter in the absence of an ORF was not sufficient to confer decompaction (Figure 6A), our results suggest that ORF transcription by Pol II is required for decompaction of the chromatin at the *GAL* locus. This is consistent with the observation in mammalian cells that only initial decompaction is mediated by transcriptional activation but that elongation by RNA polymerase is necessary for complete decompaction (Hu *et al*., 2009). We hypothesize that the removal of nucleosomes at the promoter presents a first stage of decompaction, but elongation and the concomitant eviction of nucleosomes along the ORF constitutes the major contribution to chromatin opening. This interpretation is in line with the observation that opening of the *GAL* locus is not detected when both glucose and galactose are present (Figure 4C Galactose+Glucose condition), a condition where SWI/SNF activity has been reported to evict nucleosomes at the promoter (Bryant *et al*., 2008). However, in addition to the length of the ORF, the strength of activation and possibly the composition of the 3’ and 5’ UTRs may also contribute to the level of nucleosome eviction and thus decompaction. Importantly, the absence of a decompaction response from the promoter alone also indicates that longer range interactions mediated by the promoter, for example promoter-enhancer loops, do not play a role in our system allowing us to focus on the local changes in chromatin organization. Such effects or the release from higher spatial compaction states may have contributed to the transcription-independent decompaction that was observed previously in mammalian cells (Hu *et al*., 2009).

### The role of nucleosome eviction

Histones are evicted during transcriptional elongation but repositioned after the passage of Pol II. Both eviction and deposition require the activity of histone chaperones like FACT. In highly transcribed genes like the *GAL* genes, individual transcription events follow each other closely (Lenstra *et al*., 2015). In this case, deposition of histones between rapid rounds of transcription may not be possible, and thus a net loss of histones and nucleosomes occurs. Such a response would explain the dramatic unfolding of chromatin across the *GAL* locus observed in our assay and is consistent with our observation that mutants with reduced transcriptional activity display a reduced decompaction. Here, initiation and polymerase II progression occur at lower frequencies, which could allow for more time to deposit histones after the passage of Pol II. This interpretation is also in agreement with the observed correlation between ORF length and decompaction.

FACT has previously been shown be important for nucleosome removal in the *GAL* promoter and ORF. Interestingly, the *spt16ts* mutant allowed us to at least partially uncouple transcription from decompaction: while no detectable decompaction occurred one hour after induction at the restrictive temperature in this mutant, the mRNA induction was indistinguishable from wild type cells, indicating that nucleosome and chromatin compaction are not limiting for transcription under these experimental conditions. However it is possible, that higher levels of transcription require nucleosome eviction and decompaction, since even wild type cells exhibited reduced levels of transcription at the restrictive temperature.

Combined with our modelling results, our data are consistent with the hypothesis that changes in distance are caused by nucleosome eviction. However, we cannot exclude that additional transcription-associated processes including histone tail modifications or the mRNA transcript itself also provide important contributions to the observed decompaction response.

### Outlook

Transcription initiation displays large cell-to-cell variability, and the same can be expected also for decompaction. Unfortunately, we were not able to reliably measure decompaction kinetics on a single cell level, due to the large fluctuations in distances measured over time even for cells in steady state growth conditions (Shechtman *et al*., 2017), which is also reflected in the wide distribution of distances in the population at any single time-point (Figure 1C). Therefore, further developments of reporter systems with reduced biological noise in combination with improved superresolution microscopy approaches will be needed to enable the analysis of decompaction kinetics on the single cell level. For example, the contribution of DNA sequences that are not involved in the observed response could be minimized by placing fluorescent reporters directly adjacent to the gene of interest. Adaptations of our system thus have the potential to shed further light on the dynamics of chromatin decompaction by allowing to study this process quantitatively in single cells under physiological conditions. The data from live cell microscopy experiments are highly complementary to data obtained from chromosome capture techniques and are very valuable to better understand chromatin organization and its dynamics *in vivo*. Furthermore, using CRISPR/Cas technologies, our assay could be readily transferred to higher eukaryotic systems. Optimized cassettes for the integration of LacO repeats by CRISPR/Cas have recently been created (Tasan *et al*., 2017) and the targeting of labelled CIRSPR/Cas complexes (Chen *et al*., 2013; Chen and Huang, 2014; Ma *et al*., 2016) will in the future provide a powerful and flexible tool to dissect the contribution of individual players to chromatin compaction in various model systems.

## Methods

### Plasmid construction

Plasmids were constructed using standard molecular biology techniques. Fragments generated by PCR were verified by sequencing. pKW1008 was constructed by ligating annealed oligos UC586/867 into pFA6-GFP(s65t)-kan (Longtine *et al*., 1998) using PacI/AscI to replace GFP with a FLAG-tag. pKW2695, pKW2704 and pKW3264 were constructed by ligating PCR products of primers CH4505/CH4506, CH4513/CH4514 or CH6/CH106 on genomic DNA into pKW1689 (Green *et al*., 2012) after cutting vector and insert with XhoI and SacI. pKW3035 was constructed from a PCR product of UC5681/UC5682 on pKW3010 (Backlund *et al*., 2014) cut with DraIII and ligated into pAFS135 (Straight *et al*., 1996) also cut with DraIII. pKW3681 and pKW3682 were constructed by inserting fragments generated by primers CH1211/CH1213 or CH1211/CH1177 on genomic DNA into the pFA6a backbone containing a NatMx resistance cassette (cut with PacI/XhoI). pKW3683 was constructed by inserting into the same vector a stitched PCR product of the *GAL1* promoter (CH1211/CH1218) and GST (CH1214/CH1215) with PacI/XhoI. pKW4420 was constructed in multiple steps. The final construct contains homology regions for a region on chromosome II which were amplified from genomic DNA using primers CH1795/CH1796 and CH1797/CH1798. Between the homology sites, the *GAL1* promoter amplified by CH1211/CH2043 drives a construct encoding 2xGST (CH1789/CH1794 and CH1791/CH1794) tagged with V5. It also contains the NatNT2 resistance cassette for selection and can be integrated after cutting NotI/AscI. The Gal4DBD-ER-VP16 activation construct was transferred from the construct published in (Louvion *et al*., 1993) into a Ylplac204 backbone with an additional NatMx cassette added for selection in TRP+ strains (pKW3504). All plasmids used in this study are listed in Supplementary Table 1. Primers are listed in Supplementary Table 2.

### Yeast strain construction

S. *cerevisiae* strains were constructed in the background of BY4741 and BY4742 (Brachmann *et al*., 1998) using standard yeast genetic techniques either by transformation of a linearized plasmid or of a PCR amplification product with homology to the target site (Baudin *et al*., 1993). The mutants *spt20Δ, gal80Δ, gcn5Δ, asf1Δ, snf2Δ, gal11Δ, pob3-7, spt16ts, med7-163, rpb1-1, gal4Δ, swr1Δ* and *arp6Δ* were constructed by mating of the wild type strains with the corresponding strains from the MATa, MATα deletion collections (Winzeler *et al*., 1999) or the MATα temperature sensitive collection (Li *et al*., 2011) followed by sporulation and tetrad dissection. *ADA2* was deleted by PCR-directed mutagenesis using primers CH395/CH396 on pKW1008 to generate KWY5104. Reporter strains for the experiments shown in Figure 6A were generated by transformation of KWY4067 with the PCR product of CH1272/CH1273 on pKW3681 (=> KWY6245), pKW3682 (=> KWY6247) or pKW3683 (=> KWY6250) respectively. Genotypes were confirmed by PCR. All yeast strains used in this study are listed in Supplementary Table 3.

### Yeast culture conditions

Yeast cells were cultured in complete synthetic medium with 2 % of either glucose, galactose or raffinose as indicated at 30 °C. Temperature sensitive mutants were grown at 25 °C. Some kinetic experiments with mutants showing significant growth defects were also carried out at 25 °C. All experiments were carried out with cells in exponential growth phase. For microscopy experiments, cells were usually inoculated from saturated cultures into fresh medium and grown overnight to OD 0.5-0.8 and then imaged. Alternatively, cells were diluted in the morning and grown for additional 2-3 cell cycles before the start of the experiment. Trichostatin A (Sigma Catalog #:T8552) was used at 50 μM.

### Microscopy

Cells were pre-grown in raffinose-containing medium and inoculated in 1 ml in 24 well plates overnight so that they reached an OD of 0.5-0.8 in the morning. Cells were then transferred to concanavalin A coated 384 well plates (Matrical).

3D microscopy of living cells was carried out on a SpinningDisk microscope (Yokogawa Confocal Scanner Unit CSU-W1-T2) built on a Nikon TiE body and controlled with the VisiVIEW software using dual camera acquisition mode and a 100x NA 1.49 CFI Apo TIRF objective. Cameras were Orca Flash 4.0 V2 used with 2x2 binning or iXon Ultra EMCCDs (Andor), excitation lasers were a DPSS 488 nm (200mW) and a diode 561 nm (200 mW) laser. Z scanning was performed in streaming mode with a LUDL BioPrecision2 Piezo Stage with 100 ms exposure times per frame. Filters were: Dichroic quadband DAPI/GFP/RFP/CY5, splitting filter to camera ports: 561LP, emission filters GFP/ET525/50 and mCherry ET630/75 respectively. Imaging for Figure 6A was performed on the same system in widefield-mode with excitation from Spectra X LED lines at 475 nm and 542 nm.

Time-lapse and FISH imaging experiments were carried out on a temperature controlled Nikon Ti Eclipse equipped with a Spectra X LED lamp using a Apochromat VC 100x objective NA 1.4 (Nikon) (filters: Spectra emission filters 475/28 & 542/27 and DAPI/FITC/Cy3/Cy5 Quad HC Filterset with 410/504/582/669 HC Quad dichroic and a 440/521/607/700 HC quadband filter (Semrock)) with exposure times of 50-200 ms. For timelapse experiments, induction with different sugars was carried out on stage after the first imaging timepoint by mixing with an equal volume of sugar-containing medium. Different fields of view were imaged at each timepoint to prevent phototoxicity and bleaching.

Microfluidics experiments were carried out using a CellASIC ONIX Microfluidic Platform (Merck Millipore). Cells were loaded in CellASIC ONIX plates for haploid yeast cells (Y04C) with 1 psi for 30 sec followed by short pulses (0.5 sec) of 3 psi. Medium was constantly flown across the cells at 3 psi during the entire experiment, while the medium channel that was not used was set to 0.3 psi to prevent back-flow of the other medium into the medium well. Upon medium switching, the flow rates of the two channels were reversed. The Dextran-AlexaFluor680 (Molecular Probes D34681) was added to the glucose medium at 0.1 mg/ml to monitor flow and medium switching.

### Single molecule fluorescence *in situ* hybridization

Single molecule fluorescence *in situ* hybridization was carried out according to (Mugler *et al*., 2016) with slight adaptations. The indicated strains were inoculated in synthetic medium containing 2 % raffinose and grown overnight to saturation. The next day, cells were diluted into fresh synthetic complete media containing 2 % raffinose or 2 % galactose and grown overnight at 30 °C to exponential growth phase (OD600 = 0.6-0.8). Cells were then induced by addition of glucose or galactose to final concentration of 2 % and fixed after the indicated timepoints for 15 min at 30 °C and for 15 min at 25 °C with 4 % paraformaldehyde (EM grade 32 % paraformaldehyde acqueous solution electron Microscopy Sciences 15714), washed with buffer B (1.2 M sorbitol, 100 mM KHPO_4_ at pH 7.5, 4 °C) and stored at 4 °C overnight. Cells were then spheroplasted for 20 min using 1 % 20T zymolyase in 1.2 M sorbitol, 100 mM KHPO_4_ at pH 7.5, 20 mM vanadyl ribonuclease complex and 20 μm β-mercaptoethanol, washed with buffer B to stop the spheroplasting reaction and then washed into 10 %formamide (Merck Millipore S4117) in 2x SSC.

Mixtures of DNA probes coupled to Quasar670 (Stellaris, LGC Biosearch, Novato, CA; probes were synthesized by BioCat, Heidelberg, Germany) were used for smFISH, targeting the *GAL1, GAL7* and *GAL10* ORFs (Supplementary Table 4). Per sample, 0.5 μl of each probe mix (stock 25 μM) was mixed with 2 μl of salmon-sperm DNA (10 mg ml-1, Life Technologies, 15632-011) and 2 μl yeast transfer RNA (10 mg/ml, Life Technologies, AM7119). The probe mix was denatured in 50 μl per sample of hybridization buffer F (20 % formamide, 10 mM NaHPO_4_ at pH 7.0) for 3 min at 95 °C and then mixed with 50 μl per sample hybridization buffer H (4x SSC, 4 mg/ml BSA (acetylated) and 20 mM vanadyl ribonuclease complex). Cells (approximately corresponding to the pellet of 5 ml initial culture) were resuspended in the hybridization mix and incubated for 8-12 hours at 37 °C. After four washing steps (10 % formamide/2x SSC; 0.1 % Triton/2x SSC; 2x SSC/DAPI; 2x SSC), cells were stored at 4 °C. Cells were imaged in concanavaline A coated 384 wells. Microscopy was performed with an inverted epi-fluorescence microscope (Nikon Ti) equipped with a Spectra X LED light source and a Hamamatsu Flash 4.0 sCMOS camera using a PlanApo 100 x NA 1.4 oil-immersion objective and the NIS Elements software. 31 z planes were acquired. The stack of the Qasar670 channel was acquired first due to significant bleaching. mCherry and GFP channels were acquired plane by plane to minimize shift between channels.

### Image analysis

Images were processed using FIJI (NIH ImageJ 1.51p and previous), Diatrack (Vallotton *et al*., 2017) and MATLAB^®^ (MathWorks^®^). For 2D analysis of distances on single image planes (for all kinetic experiments shown), positions of chromosome locations marked with LacI or TetR were detected by a 2D Gaussian fit using custom written scripts in MATLAB^®^ (MathWorks^®^) (Dultz *et al*., 2016). Distances between detected spots were calculated. Due to negligible shift between the two channels in these datasets, correction was not required. For 3D analysis of distances (data acquired on dual camera spinning disk microscope), positions of chromosome location marked with LacI or TetR were detected in Diatrack in 3D using across-color tracking to pair corresponding spots from the same cell (parameters used: ‘exclude blurred’ 0.07-0.2, ‘exclude dim’ 20-200, full width half maximum for Gaussian fitting 2.5). Subsequently, positions were corrected for shift between the two channels using bead images (1 μm TetraSpeck fluorescent microspheres, ThermoFisher Scientific) acquired under identical imaging conditions. For shift correction, the field of view was subdivided into 150x150 pixel large squares and the mean shift for beads acquired in this region was applied to correct the positions of chromosome locations acquired in this region of the camera. The distances between corrected positions were calculated and used for further analysis. For FISH analysis, positions of chromosome locations marked with LacI or TetR were detected in Diatrack in 3D using across-color tracking to pair corresponding spots from the same cell. Positions of transcriptions spots were detected on maximum intensity projections of the Qasar670 channel in Diatrack (parameters: remove dim 200, remove blurred 0.09-0.1, high precision fitting 2.5) and used to categorize cells as having or not having a transcription spot for separate analysis of the two populations. For analysis of transcription spot intensities, transcription spots were detected on 3D stacks in Diatrack and intensity values were obtained from the Diatrack session file. Only the intensity of those transcription spots for which corresponding chromosome locations marked with LacI or TetR could be detected in Diatrack were included in the analysis. Bootstrapping was used to obtain 95 % confidence intervals of the medians and mean. Boxplots were plotted in MATLAB^®^ using the function boxplot with the default variables: the central line corresponds to the median, the boxes extend from the 25^th^ to the 75^th^ percentile and outliers are defined as data points greater than q3 + 1.5 × (q3 – q1) or less than q1 – 1.5 × (q3 – q1). q1 and q3 are the 25th and 75th percentiles of the sample data, respectively. The whisker in each case extends to the most extreme data value that is not an outlier. All analysis code is available upon request.

### Quantitative real-time PCR

qPCR was performed as described previously (Dultz *et al*., 2016): 1 ml of cells at OD600 0.8-1 was harvested by centrifugation and snap frozen in liquid nitrogen. RNA was extracted using the RNeasy kit from Qiagen via mechanical disruption. 300 ng of total extracted RNA was used for reverse transcription. The RNA was first treated with DNase I using the DNA-free kit from Ambion according to the protocol of the manufacturer. Reverse transcription was performed according to the protocol of the manufacturer using Superscript II reverse transcriptase (Invitrogen) with random hexamer primers. Quantitative Real Time PCR was performed on a StepOnePlus Instrument (Invitrogen) using Absolute Blue QPCR Mix with SYBR Green and ROX (ThermoFisher Scientific). All experiments were carried out in three technical replicates and three biological replicates. Data was analyzed by the comparative CT method using *ACT1* as endogenous control. Primers used for qPCR are listed in Supplementary Table 2.

### Wester blotting

For Western blotting, cells were grown in the presence of plain synthetic medium with 2% raffinose or 2% galactose (supplemented as indicated with 1% DMSO or 1% DMSO + 50 uM TSA) for 16 hours to log phase and harvested by centrifugation. Cells were lysed with 0.1M NaOH and resuspended in reducing sample buffer. The equivalent of 1-2 OD of culture was separated on a 15% SDS-PAGE, blotted onto nitrocellulose membrane and labelled with an antibody against anti-acetylated proteins (Cell Signaling #9441) at 1:1000 dilution and anti-Fibrillarin antibody (ThermoFisher Scientific 38F3) at 1:2000 dilution. Secondary antibodies were goat anti mouse-Alexa680 (ThermoFisher Scientific A-21057) used at 1:10 000 dilution and goat anti rabbit -IRDye800 (Rockland) used at 1:5000 dilution. Staining was visualized on an Odyssey CLx Infrared Imaging System (Li-Cor).

### Statistical analysis

The effects of treatment and strain on the median distance value in 3D measurements or on fold induction in gene expression measurements were quantified and statistically tested using a linear mixed effect model with strain and treatment as fixed effects and the experimental day as random effect. Function lmer() of package lme4 of program R was used (Bates *et al*., 2015; Team, 2017). Reported corrected p-values were computed by post-hoc tests using function glht() of package multcomp of program R (Hothorn *et al*., 2008).

### Computational modeling

Average spatial distances were calculated from ensembles containing 1000 structure models of the entire haploid yeast genome. Ensembles were generated for each of the two *GAL* locus activation states (active/galactose and inactive/glucose), as described in Tjong *et al*., 2012 and Dultz *et al*., 2016. All chromosomes were modeled as chains of connecting beads subject to a number of spatial restraints. In particular, all chromosomes were confined inside a nucleus with a radius of 1 μm; the nucleolus and spindle pole body are placed on opposite sides of the nucleus along the central nuclear axis; all non-ribosomal-DNA gene regions are excluded from the nucleolar volume; centromeric regions are proximal to the spindle pole body, whereas telomeric regions are tethered to the nuclear envelope (allowing a maximal distance between telomeres and the nuclear envelope of 50 nm). Each bead accommodated ~3.2kb of genome sequence as described in (Tjong *et al*., 2012).

To reproduce the experimental data, we modeled a gradual decrease in chromatin compaction in the proximity of the *GAL* locus. For the 60 kb region starting at the position of the *GAL* locus, the compaction ratio was set to 1.6 kb per bead. We then determined the optimal chromatin compaction for the 6.2 kb *GAL* locus so that the models reproduce most closely the experimentally observed 3D fluorophore distances in both activation states. The *GAL* locus compaction in the inactive glucose state was found to be ~375 bp/bead whereas in the active galactose state it was set to ~200 bp/bead.

Each ensemble of genome structures was generated from 1000 independent simulations, each starting from random configurations. The optimization procedure consisted of a simulated annealing Molecular Dynamics run followed by Conjugate Gradients score minimization, both performed using IMP (Russel *et al*., 2012).

## Author contributions

Conception and design of the work: ED and KW; Data collection: ED and RM; Data analysis and interpretation: ED and RM; development of code for image analysis: ED and PV; modelling: GP & FA Drafting the article: ED; critical revision of the article: ED, RM, PV, GP, FA & KW.

## Acknowledgements

We are grateful to R. Sachdev, A. Kralt, S. Heinrich and other members of the Weis laboratory for critical discussions and suggestions on the manuscript. We thank Joachim Hehl from the Scientific Center for Optical and Electron Microscopy of ETHZ (ScopeM) for support with microscopy and Andreas Steingötter from the Statistical Consulting Group at ETH Zurich for help with statistical analysis. This work was supported by the Swiss National Fond (SNF 159731) and NIH (U54DK107981 to F.A).

## Abbreviations

TSA: trichostatin A
ORF: open reading frame
qPCR: quantitative PCR
SE: standard error
GST: Glutathione S transferase
smFISH: single molecule fluorescence *in situ* hybridization
Pol II: RNA polymerase II

---

**Supplemental Figure 1:**
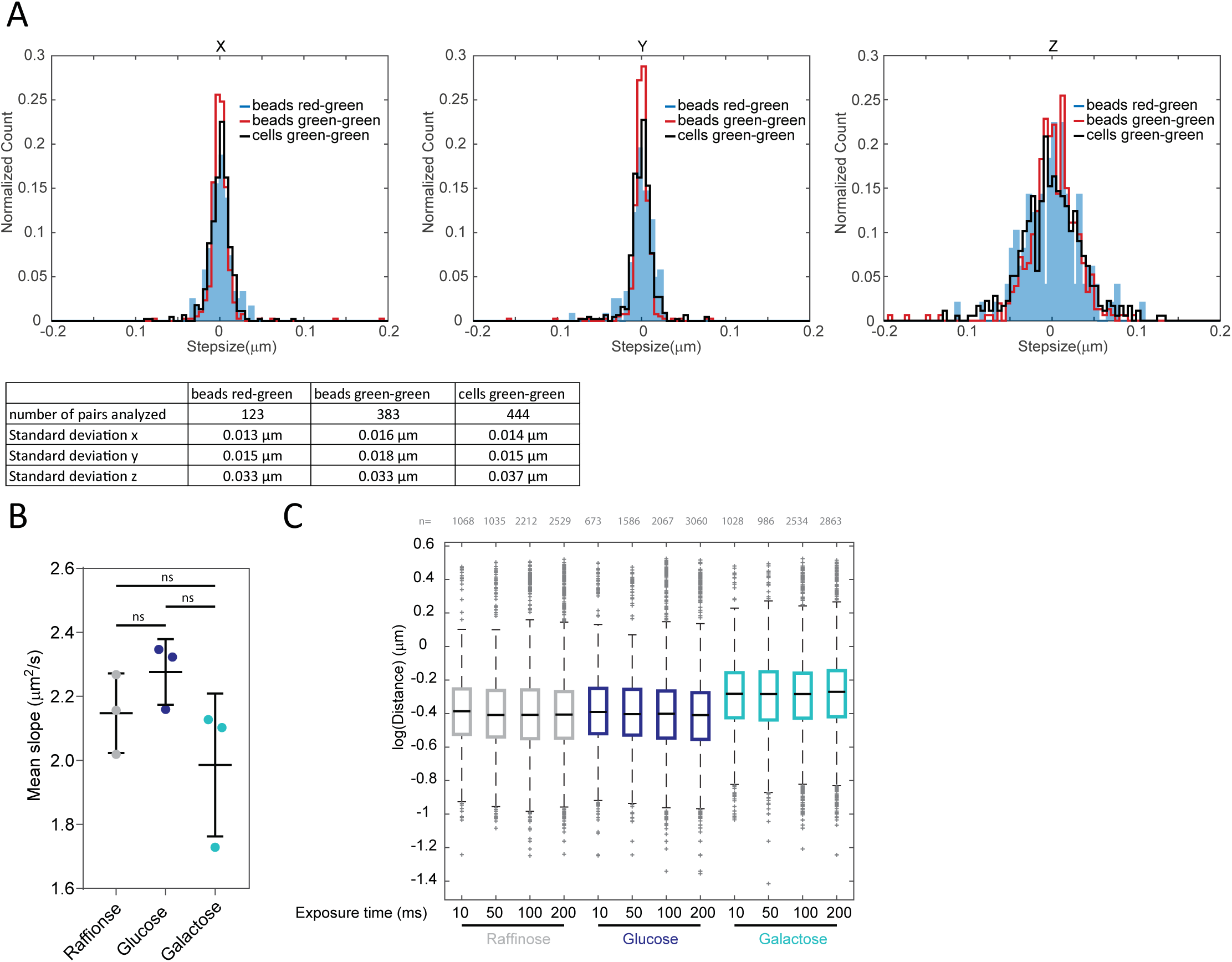
Measurement precision. (A) Precision of distance measurements was assessed using beads and cells. Stacks of 1 mm fluorescent microspheres were acquired with small shifts of the stage (1-2.5 μm in x and y, 300-400 nm in z) with identical imaging settings used for imaging chromosomal loci in 3D experiments throughout this work. The distance between beads in a single channel (green-green) was analyzed to determine measurement precision. The standard deviation should converge to 0, since the stage moved all beads by the identical distance. The measured distances (normalized by the mean shift) are shown in histograms. The same procedure was applied to fixed cells to assess the contribution of non-perfect biological signals. To assess the additional error introduced by the correction for shift between cameras, the same analysis was performed after application of the shift correction procedure to bead images acquired in the red channel compared to bead images acquired in the green channel after stage shift. (B) Movies of TetR-mCherry-labelleled chromosomal locus in the wild type reporter strain were acquired at 100 ms resolution using triggered acquisition mode. Movement of GFP spots was tracked in Diatrack and mean square displacement (msd) curves were calculated for each track. The first second of each msd-curve was fit with a linear regression in MATLAB using polyfit. The mean slope of three biological replicates is shown (lines represent means and standard deviations; number of tracks in each replicate: raffinose: 283,143,182; glucose: 272, 123, 195; galactose: 275, 111, 223). Adjusted p-values were obtained by applying an ordinary one-way ANOVA and Sidak’s multiple comparisons test in Prism. Note that possible reduced mobility in galactose compared to glucose (although not significant) would lead to a decrease rather than an increase in the observed distance. (C) 3D distance datasets acquired at different exposure times. A Kruskal-Wallis test and Dunn’s multiple comparison correction in Prism showed no significant differences (p>0.05) in the distributions of different exposure times (each exposure times was tested against 100 ms, which is the exposure time used for all 3D experiments throughout the paper). Shown is one of three biological replicates as a boxplot (Red lines show the median, boxes extend to the 25th and 75th percentile. Red crosses represent outliers, which are defined as points that are greater than q3 + w × (q3 – q1) or less than q1 – w × (q3 – q1), where w is the maximum whisker length, and q1 and q3 are the 25th and 75th percentiles of the sample data, respectively.).

**Supplemental Figure 2:**
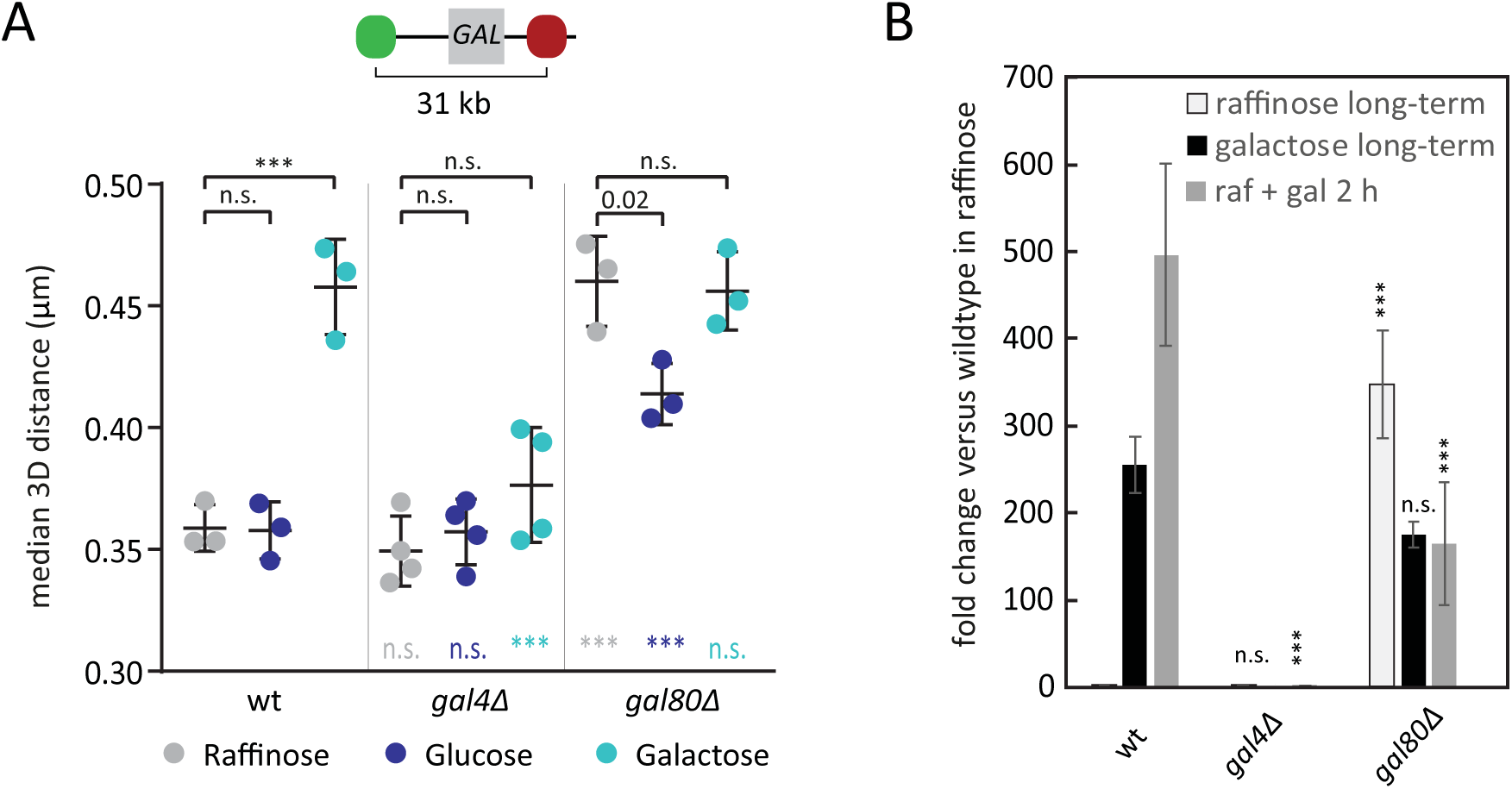
Transcription activation mutants also affect opening of the *GAL* locus. (A) The median distance between LacO and TetO repeats in cell populations one hour after addition of the indicated carbon source to the raffinose containing culture medium to a final concentration of 2% (raffinose 4%). Each dot represents the median of an independent experiment with at least 100 analyzed cells (typically 600-1000). Lines and error bars indicate means and standard deviations. *** adjusted p-value < 0.001, n.s. not significant (p > 0.05). Colored p-values represent the adjusted p-values of a comparison to the wild type in the same condition. (B) Fold induction of *GAL10* mRNA compared to wild type cells grown in raffinose determined by quantitative PCR (normalized *for ACT1* mRNA as endogenous control). Means of 2-4 biological replicates are shown. Error bars represent standard errors.

**Supplemental Figure 3:**
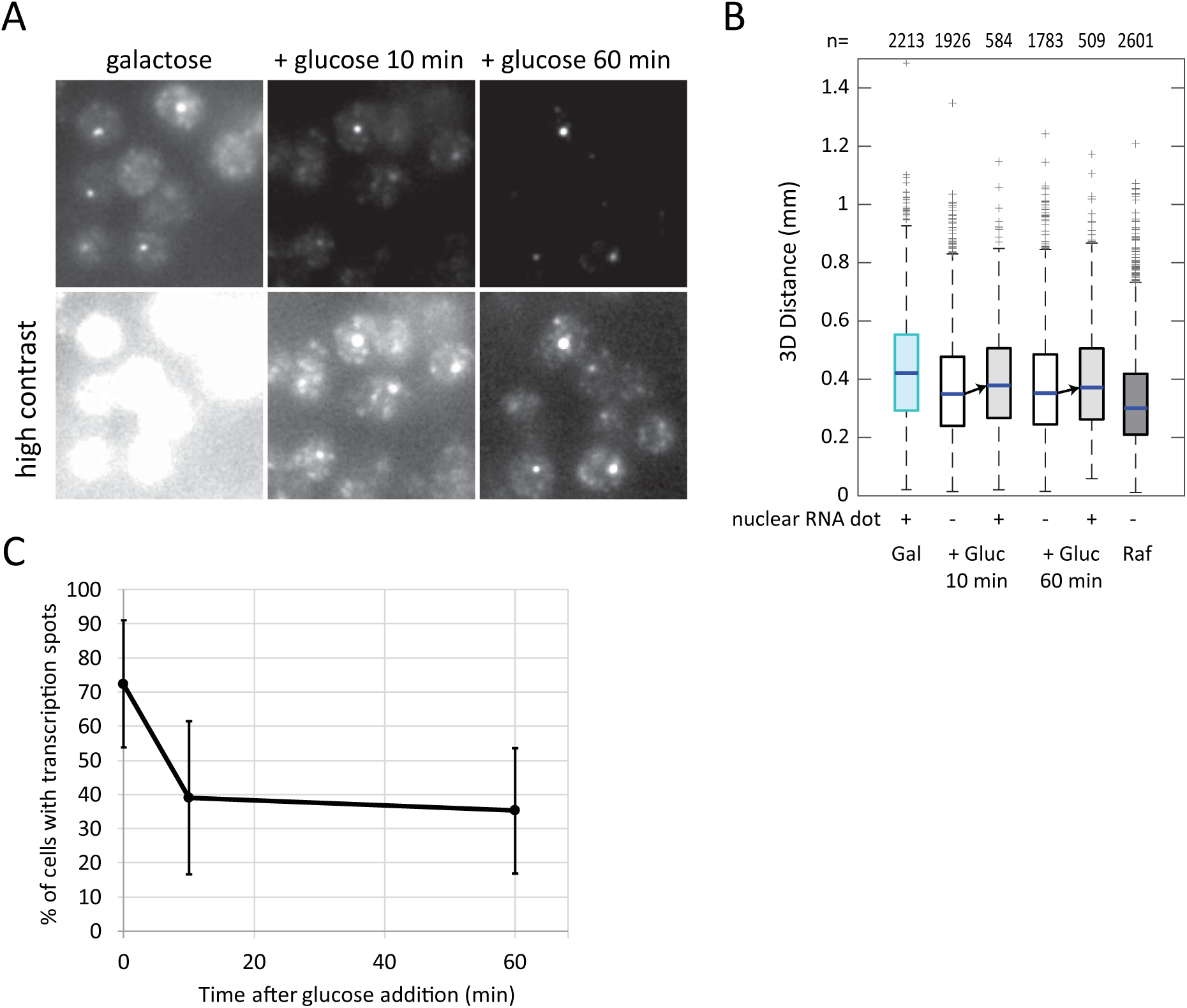
RNA dot persists after glucose repression and correlates with more open chromatin. (A) Maximum projections of smFISH for *GAL1, GAL7* and *GAL10* mRNAs detected by oligos labelled with Qasar670 in cells grown to exponential phase in galactose and then repressed by the addition of glucose to a final concentration of 2%. Images from all conditions are shown at identical contrast settings. Bottom panels show the same images with higher contrast. Images were filtered with a Gaussian filter for presentation purposes. Scale bar represents 2 μm. (B) Boxplot showing the distance distribution in cells grown either in galactose or raffinose continuously or repressed by addition of glucose to galactose grown cells. Distributions of cells with or without transcription spot are analyzed separately 10 and 60 min after glucose repression. Data from one representative experiment out of two biological replicates is shown. n: number of cells analyzed. (C) Mean % of cells with nuclear RNA dot from two biological replicates (error bars represent standard deviation).

**Supplemental Figure 4:**
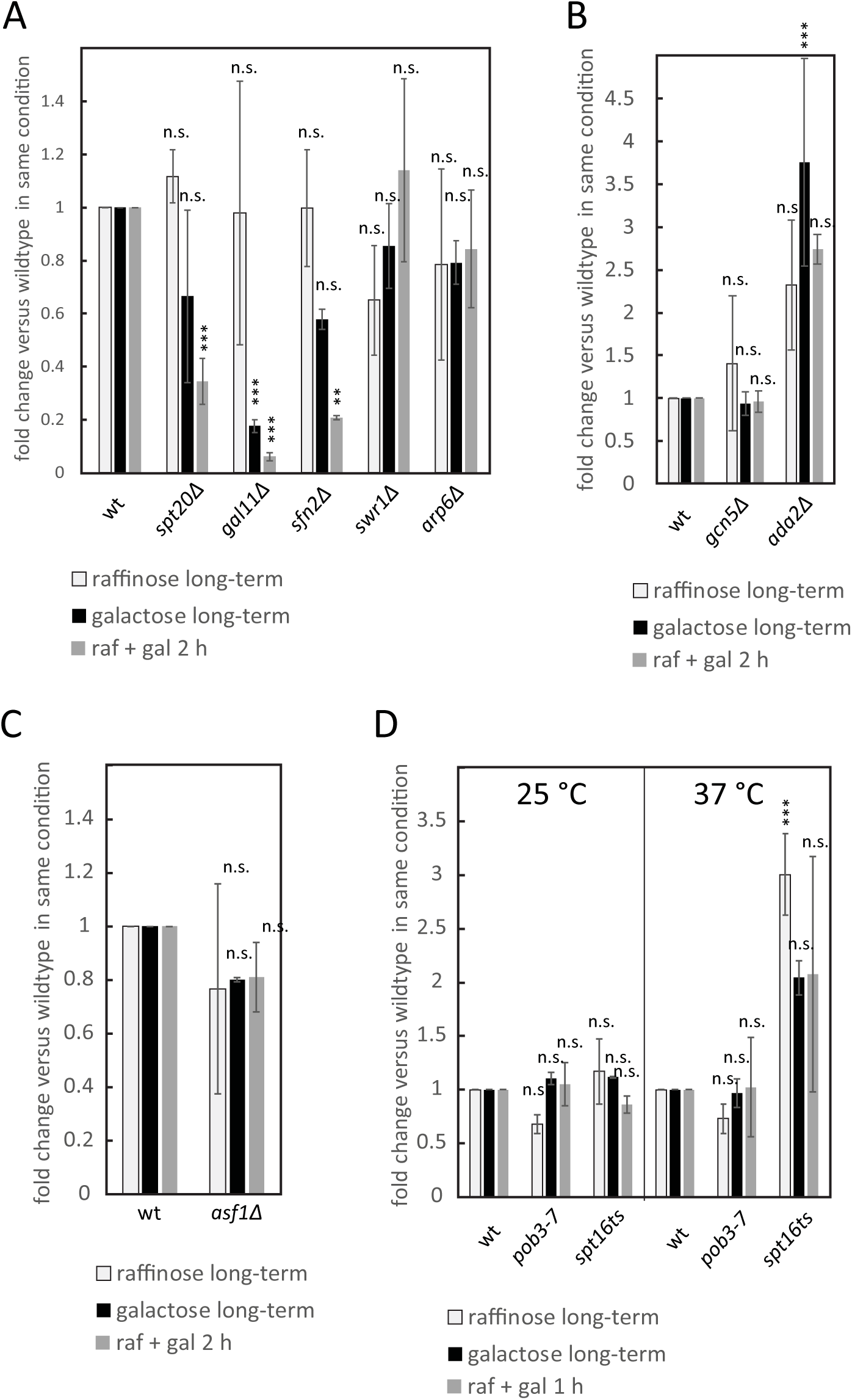
Gene expression data for the mutants analyzed in the manuscript normalized by the level of *GAL10* mRNA present in the corresponding wild type strain in the same condition. Data are the same as plotted in the manuscript with different normalization. Each bar represents the mean of 3-4 biological replicates, the error bars represent the standard error. Differences between wild type and mutants were tested in a mixed linear model with multiple comparison post-hoc tests (compare Material and Methods). Adjusted p-values: *** p< 0.001, ** p<0.01, * p< 0.05, n.s. p>0.05.

**Supplemental Figure 5:**
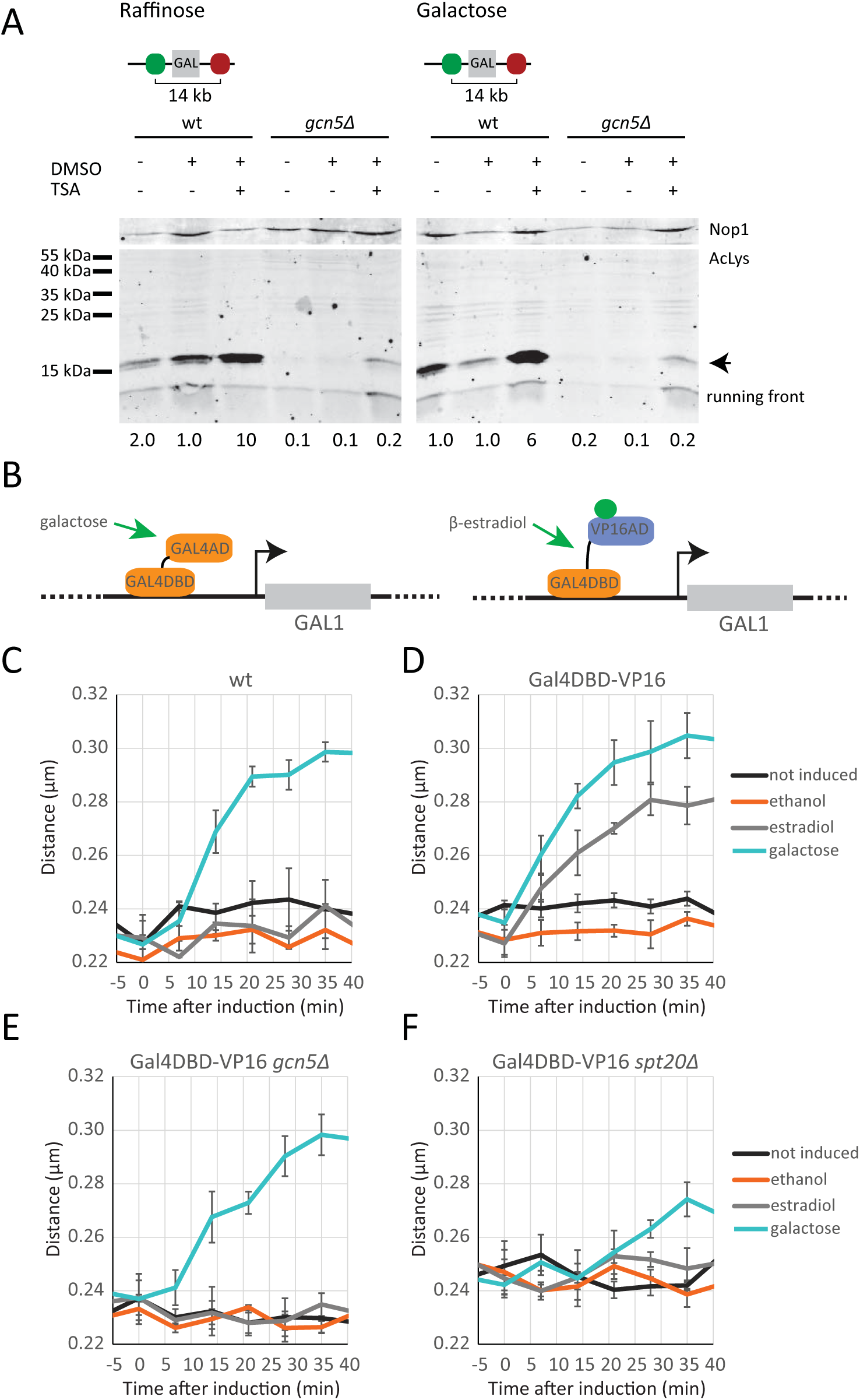
Decompaction mediated by VP16 depends on SAGA’s HAT activity. (A) Western blot showing increased levels of lysine acetylation upon TSA treatment in proteins in the size range of histones (arrow). Quantification indicates levels normalized to Nop1 signal and the DMSO treated wild type sample. Shown is one of three biological replicates. (B) Schematic representation of the VP16 induction system. The Gal4 DNA binding domain (Gal4DBD) is fused to the activation domain of VP16 linked by the hormone binding domain of human estrogen receptor. Upon addition of β-estradiol to the cells, the fusion protein is released from its interaction with a heat shock protein and can activate transcription from promoters containing a Gal4 recognition motif. (C-F) Decompaction kinetics in response to activation via galactose or β-estradiol. Cells were grown to exponential phase in medium with 2 % raffinose. Induction was carried out on the microscope stage at timepoint 0 with galactose (2 % final concentration) or β-estradiol (1mM final concentration). Shown are the means of three biological replicates with the standard error represented by the error bars. (C) Wild type. (D) Wild type expressing β-estradiol responsive VP16 activator. (E) *gcn5Δ* expressing β-estradiol responsive VP16 activator. (F) *spt20Δ* expressing β-estradiol responsive VP16 activator.

**Supplemental Figure 6:**
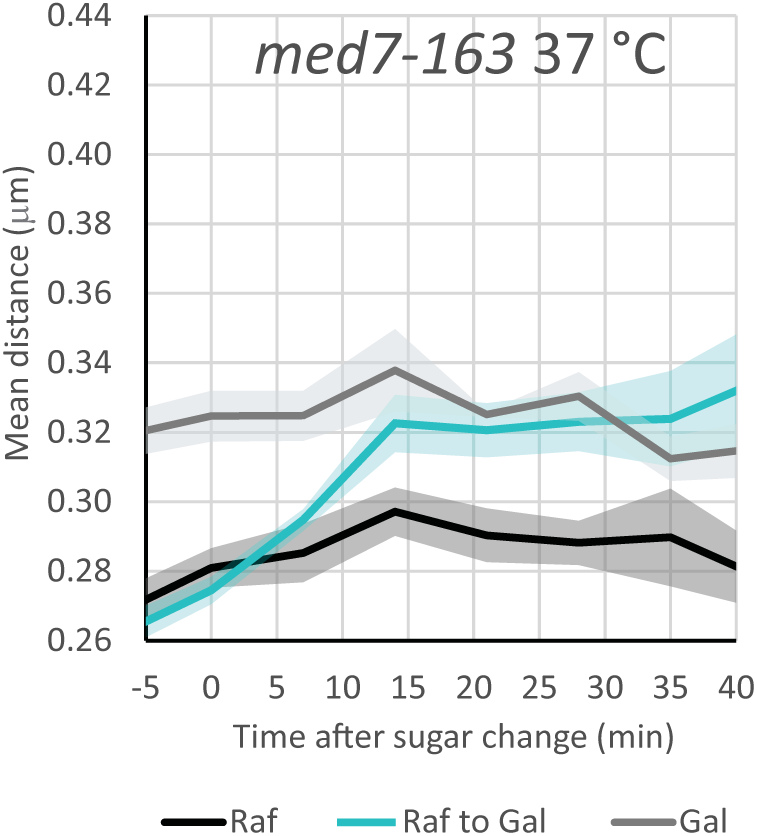
Kinetics of opening in *med7-163* temperature sensitive mutant at restrictive temperatures. Cells were grown overnight in medium with 2 % raffinose or 2 % galactose at 25 °C and shifted to 37 °C one hour before timepoint 0. Galactose was added or not added to 2 % final concentration at timepoint 0. Shown are the means of three biological replicates with the standard error represented by the shaded areas.

**Supplemental Figure 7:**
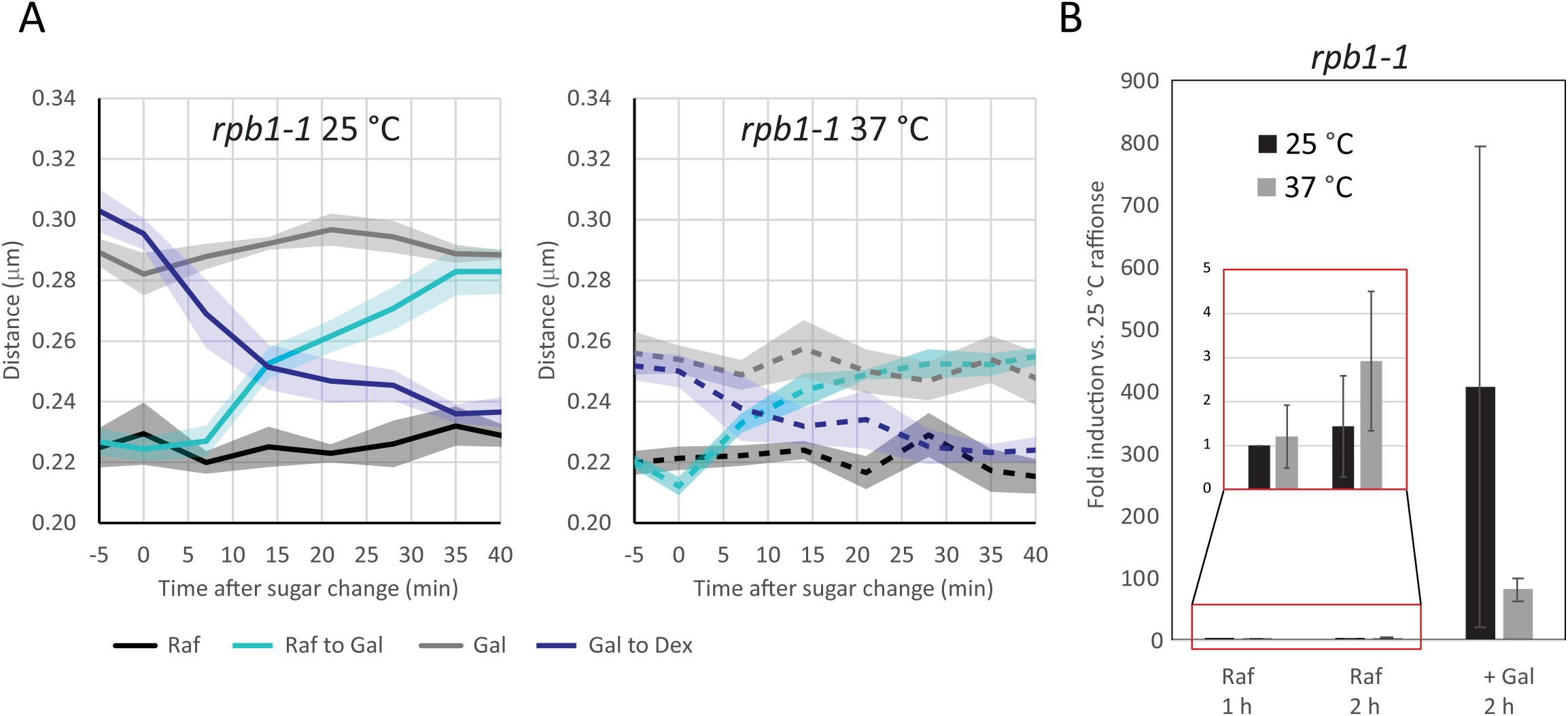
Transcription correlates with opening. (A) Kinetics of opening and closing in *rpb1-1* temperature sensitive mutant at permissive and restrictive temperatures. Cells were grown overnight in medium with 2 % raffinose or 2 % galactose at 25 °C and shifted to 37 °C were indicated one hour before timepoint 0. Galactose or glucose respectively were added or not added to 2 % final concentration at timepoint 0. Shown are the means of three biological replicates with the standard error represented by the shaded areas. (B) Fold expression of *GAL10* mRNA relative to growth at 25 °C in raffinose (normalized for *ACT1* mRNA as endogenous control). Cells were grown in 2 % raffinose into exponential phase. Cells were then shifted to the indicated temperature for one hour and subsequently induced with galactose (+gal) or not for one more hour. Means of three biological replicates are shown. Error bars represent standard errors. Inset shows blow-up for uninduced (raffinose) growth.

